# Fermentation of the Edible Brown Seaweed *Alaria esculenta* by *Lactiplantibacillus plantarum* affects nutritional profile and the content of potentially toxic elements

**DOI:** 10.64898/2026.05.05.723112

**Authors:** Supansa Westman, Thamani Freedom Gondo, Madeleine Jönsson, Maren Sæther, Jon Funderud, Wender Bredie, Lilia Ahrné, Olof Böök, Dragana Stanojević, Anne Elsser-Gravesen, Charlotta Turner, Eva Nordberg Karlsson

## Abstract

Edible seaweed has the potential to become a valuable marine resource for food applications due to its potential health benefits and ecological sustainability. The brown seaweed *Alaria esculenta* is rich in essential minerals, vitamins, and dietary fibers, making it a nutritious food source. Fermentation, as a traditional preservation method, can enhance seaweed shelf-life and be useful for the development of new foods/ beverages. In this study, the effects of fermentation of *A. esculenta,* by the lactic acid bacterium (LAB) *Lactiplantibacillus plantarum,* on the nutritional profile, and the content of potentially toxic elements, was investigated. *L. plantarum* was successfully cultivated on *A. esculenta* using two modes of operation, submerged (SmF) and solid-state fermentation (SSF), resulting in production of cells and lactic acid, and reduction of the pH to below 4.3 within 3 days, which was not achieved in parallel spontaneous fermentations using indigenous seaweed microbiota. *A. esculenta*‘s macro-nutritional profile was altered, reducing mannitol but increasing fucose and glucose content (after acid hydrolysis) while also concentrating the protein content. LAB fermentation significantly increased the concentration of antioxidant phenolic compounds, such as phloroglucinol, syringic acid, and epicatechin, compared to untreated samples. However, lipophilic compounds like carotenoids decreased after both spontaneous and LAB-fermentation. A reduction in total mineral content was observed after LAB fermentation and water soaking, and SmF with *L. plantarum* effectively reduced arsenic and iodine levels. Overall, fermentation using *L. plantarum* showed potential as a bio-preservation method for the edible brown seaweed, *A. esculenta,* improving its nutritional profile and enhancing food safety.

## 1 Introduction

Edible seaweeds, or macroalgae, can become important marine resources for food applications as they are considered healthy and ecologically sustainable (1). Seaweeds grow quickly without the need for arable land or freshwater, and are rich in nutrients and bioactive compounds, beneficial to human health. Despite an increasing seaweed production worldwide, exploitation for human diets remains underdeveloped in Western societies. Barriers for its introduction into new markets include unfamiliarity, unavailability, and lack of knowledge on efficient stabilization techniques (2). Currently, techniques to avoid rapid postharvest deterioration (due to e.g. high moisture), include drying, storage at low temperatures (e.g. freezing), pretreatment at elevated temperature (blanching), and more recently also preservation via fermentation, where the latter not only leads to extended shelf-life, but also to improved nutritional values (3). Consumer interest in fermented foods is also increasing due to potential nutrient and health benefits (3).

Fermentation is a traditional method for preserving perishable raw materials and can be useful for development of new foods/ beverages, such as new seaweed-based products. During fermentation, bioactive compounds, *e.g.* organic acids, phenolic compounds, amino acids, and volatile fatty acids are released from microorganisms, contributing to unique sensory and functional properties of the product. For seaweeds, fermentation for food applications has recently been developed (4). While several microorganisms have been used, fermentation by lactic acid bacteria (LABs) has been a primary choice. LABs are non-spore forming, gram-positive bacteria whose main product of sugar fermentation is lactic acid. The production of lactic acid throughout the process, lowers pH, and inhibits growth of spoilage and pathogenic bacteria, both via competition for nutrients, and by the production of organic acid (lactic acid, and dependent on strain additional metabolites e.g. ethanol, hydrogen peroxide, and bacteriocins), resulting in bio-preservation (3). Moreover, some LABs have been recognized as probiotics, and incorporated in products for health-promoting properties that can protect against certain human and animal diseases (5, 6).

Solid-state fermentation (SSF) and submerged fermentation (SmF) are two predominant fermentation techniques, each with unique advantages and applications. SSF involves the cultivation of microorganisms on solid materials with minimal free water, making it effective for processes involving filamentous fungi and substrates like agricultural residues (7). This method often leads to higher yields of secondary metabolites, but it presents challenges for process control and scalability. According to Viniegra-Gonzalez et al. (8), SSF processes are advantageous for products that require low water activity and high aeration. On the other hand, SmF entails growing microorganisms in a liquid medium, which allows for a more precise control of environmental conditions, making it ideal for bacterial cultures and primary metabolite production. SmF is extensively used in industrial applications for producing antibiotics, enzymes, and organic acids due to its ease of scalability and process optimization. Thomas et al.(9) emphasized that SmF provides a homogeneous environment that facilitates efficient distribution of nutrients and oxygen. The choice between SSF and SmF is influenced by factors such as the type of microorganism, the nature of the product, and the specific requirements of the fermentation process. A review by Pandey et al.(10) notes that the selection of fermentation strategy is crucial and depends on the physicochemical characteristics of the substrate and the economic feasibility of the process.

Kelps, including *Alaria esculenta*, are abundant in dietary fiber, which comprises up to 70% of their dry matter. This fiber fraction includes both water-soluble and insoluble structural, storage, and functional polysaccharides, many of which exhibit bioactive or prebiotic properties (14). The protein content is typically up to 20 % of the dry weight. Additionally, seaweeds are sources of vitamins, polyphenols (*e.g.,* antioxidants), minerals, and other bioactive molecules (11, 12). Kelps also contain mannitol as a fermentable free sugar alcohol available as substrate for certain microorganisms, including *Lactiplantibacillus plantarum,* which has been reported to utilize seaweed carbohydrates in fermentation (16), and is a LAB-choice for many plant-based fermented products (6, 13). During the fermentation process, mannitol is converted into organic acids, which are beneficial from both storage and health perspectives. Previous studies have shown enhancement of the sensory characteristics of kelp after lactic acid fermentation, with demonstrated alteration of the aroma compound profile (14), less salty taste, and reduced slimy look (15). However, more studies on the effect of lactic acid fermentation on seaweed quality concerning nutritional value, sensory characteristics, and food safety are needed.

From the food safety perspective, concerns have emerged regarding the content of potentially toxic elements (PTEs), such as arsenic, cadmium, mercury, lead, and iodine (18). Although food safety regulations for seaweed in Western countries are still fragmented, the French Agency for Food, Environmental and Occupational Health & Safety (ANSES) (16) and the Algae Technology & Information Centre (CEVA)(17) have issued maximum permissible levels for certain elements in seaweed. Moreover, the European Food Safety Authority (EFSA) has established tolerable daily/weekly intake levels, and the European Commission has specified maximum limits for various contaminants in food products (18). As some seaweed harvests have been shown to exceed PTE-limits proposed by ANSES or CEVA, efforts are made to reduce the PTE content (18), and for this purpose, fermentation as a processing method also seems promising. Bruhn et al. (15) could, for instance, reduce the content of cadmium (− 35%) and mercury (− 37%) in sugar kelp after fermentation.

In this study, the primary goal was to examine how submerged (SmF) and solid-state (SSF) fermentation methods by *L. plantarum* impact the quality of the brown seaweed *A. esculenta* concerning its nutritional value as well as the food safety. The assessment of nutritional value after fermentation included analysis of protein, antioxidants, including phenolic compounds, and minerals, and food safety parameters included monitoring of PTEs. Improving the preservative effects on seaweed can help to increase the usability of this perishable marine food source.

## 2 Materials and Methods

### 2.1 Seaweed raw materials

Brown seaweed, *Alaria esculenta* (L.) Grev., harvested in late April by Seaweed Solutions A/S, Norway, was used as substrate for seaweed fermentation experiments. After harvesting, the fresh seaweed was frozen immediately, transferred to the laboratory, and stored in the freezer (−20 °C). Frozen fresh seaweed was weighed and cut with a meat mincer (Menuett, Jula AB, Sweden) while it was still frozen to prevent the loss of fermentable compounds during the thawing process. Frozen minced seaweeds were immediately used for further treatments.

### 2.2 Lactic acid bacteria and pre-culture preparation

*Lactiplantibacillus plantarum* (Orla-Jensen 1919) Zheng et al. 2020 was obtained from Aventure AB (Lund, Sweden) and was stored at −80 °C. Pre-cultures were prepared based on the recommendations obtained from the company. In brief, 0.1 g freeze-dried bacteria, estimated equal to 10^10^ CFU, was suspended in 10 mL of 0.9 % NaCl solution to reach an approximate cell concentration of 10^9^ CFU/mL. The pre-cultures were vortexed for 2 min and were thereafter left to rest, allowing the freeze-dried bacterial powder to rehydrate under sterile conditions at room temperature for 30 min. Rehydrated bacteria were vortexed and used for inoculation.

### 2.3 Chemicals and media

De Man-Rogosa-Sharpe (MRS) agar and MRS broth were used as cultivation media. MRS broth was prepared using peptone from casein (10 g/L), meat extract (8 g/L), yeast extract (4 g/L), di-potassium hydrogen phosphate (2 g/L), di-ammonium hydrogen citrate (2 g/L), sodium acetate (5 g/L), magnesium sulphate heptahydrate (0.2 g/L), manganese sulphate monohydrate (0.04 g/L), and twee 80 (1 g/L). In addition, stock solutions of glucose (100 g/L) and mannitol (100 g/L) were prepared separately for single-substrate fermentations to evaluate the performance of the bacterium on carbohydrate sources commonly present in brown seaweed. A 0.9 % NaCl solution was also prepared for pre-inoculum and cell cultivation. All chemical solutions were autoclaved at 121 °C for 20 min.

Chemicals used for the preparation of MRS medium, stock solutions, and standards were obtained from Sigma life sciences and/or Sigma Aldrich. Milli-Q water was purified with a 0.2 μm filter.

### 2.4 Lactic acid fermentation processes

#### 2.4.1 Fermentation in defined MRS media with single substrates

An experiment was designed to demonstrate the growth of *L. plantarum* on the monosaccharide glucose or the sugar alcohol mannitol, available in brown seaweed biomass (Fig. 1). One mL of pre-culture (section 2.2) was inoculated into serum glass bottles containing 100 mL of MRS medium with either 30 g/L of mannitol, 30 g/L of glucose (positive control), or water (negative control) as carbohydrate source (Section 2.3), and closed with butyl rubber seals and aluminum caps. Bacterial cultivations were performed in an incubator at 37 °C for 24 h under semi-anaerobic conditions. The experiment was performed in triplicate, and the results are presented as mean ± standard deviation.

**Figure 1.**
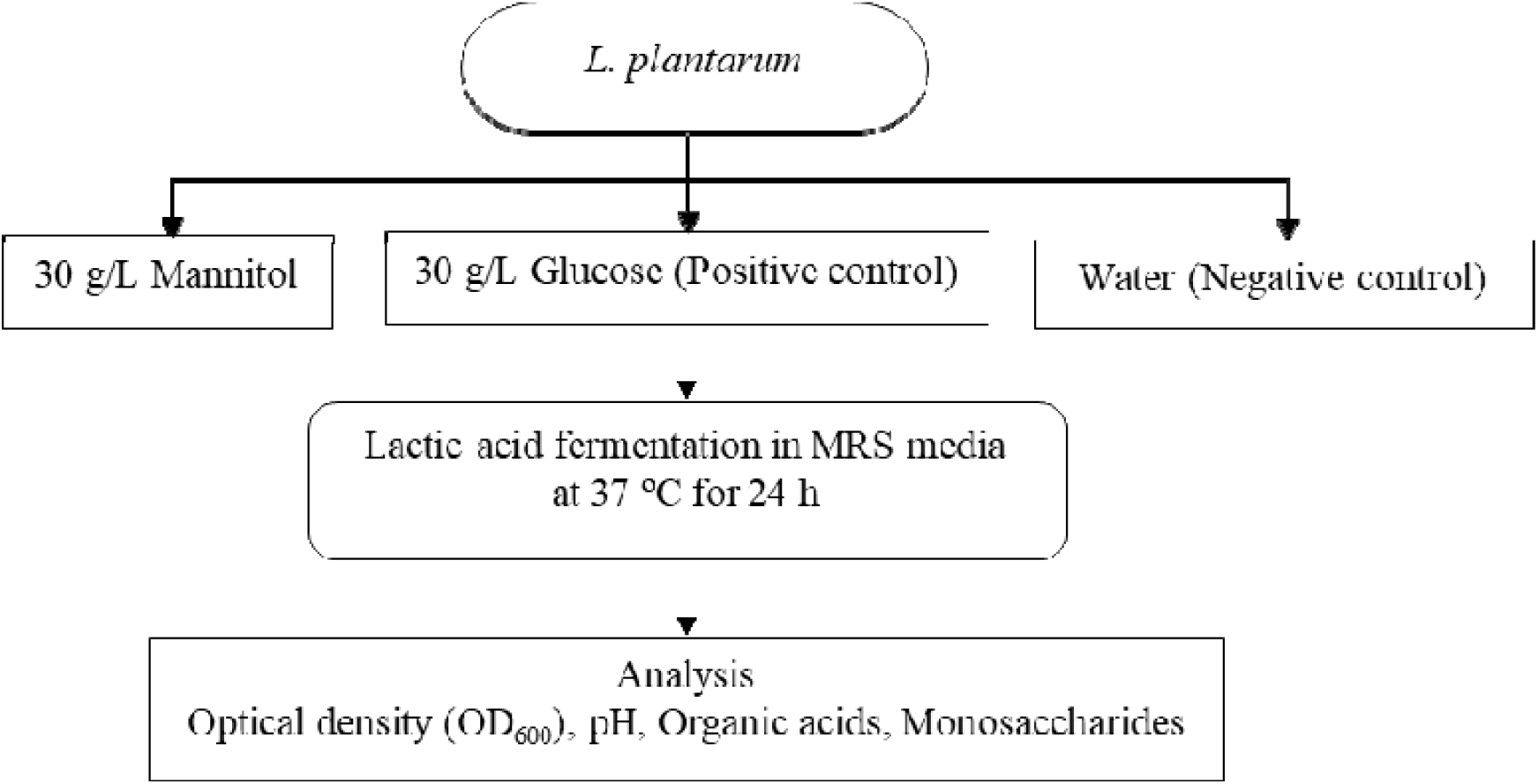
Flow chart illustrating the fermentation design of *Lactiplantibacillus plantarum* on seaweed-relevant carbohydrate sources.

#### 2.4.2 Fermentation of the seaweed substrate

Submerged lactic acid fermentation (SmF-LA) and solid-state lactic acid fermentation (SSF-LA) of ground *A. esculenta* (AE) were performed according to the following recipes:

For SmF-LA, frozen ground AE (Section 2.1) with a final concentration of 60 % w/v was mixed with 40 % v/v of Milli-Q water to make a slurry, which was aliquoted in 120 mL sterile glass bottles (Schott, Germany) and closed with screw caps. Bottles containing the cold seaweed slurry were placed in an incubator at 37 °C until the temperature of the slurry reached 30 °C, and were subsequently inoculated with pre-cultures (section 2.2) to reach an approximate final cell concentration of 5×10^7^ CFU/mL. Lactic acid fermentations were carried out in batch mode under semi-anaerobic conditions at 37 °C for 72 h. After fermentation, the seaweed solid fraction was separated from the liquid part by centrifugation at 64000 xg for 30 min, rinsed with 500 mL of Milli-Q water, and again centrifuged under the same conditions. Samples were then lyophilized, ground, and sieved (2 mm mesh) and were stored at −20 °C until further analysis. The fermentations were performed in triplicate, and the results were presented as mean ± standard deviation. Spontaneous fermentations without adding bacteria (i.e., negative control; SmF-SP) were also performed under otherwise identical conditions.

Inoculated solid-state lactic acid fermentation (SSF-LA) and spontaneous solid-state fermentation (negative control; SSF-SP) of AE were set up under identical conditions as for the submerged fermentation, but without adding water. A flow chart representation of the seaweed fermentation processes is presented in Fig. 2. The fermentations were performed in triplicate, and the results were presented as mean ± standard deviation.

**Figure 2.**
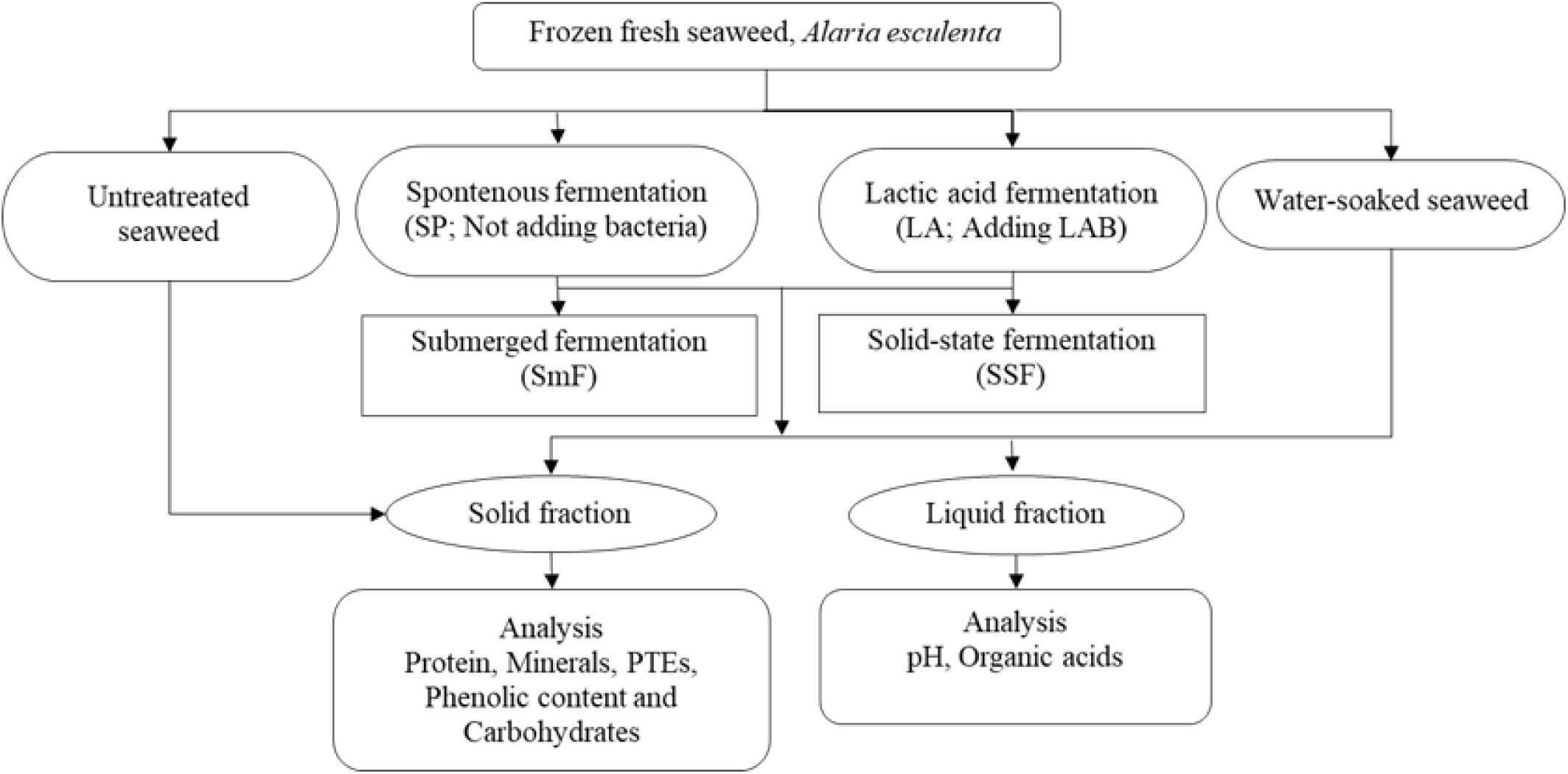
Flow chart representation of the seaweed fermentation processes.

### 2.5 Water-soaking

Soaked seaweed was prepared under identical conditions as for SmF-LA before inoculation of *L. plantarum.* This treatment was used as the negative control for SmF-LA to rule out the effect of soaking on subsequent seaweed qualities. Cold seaweed samples from Section 2.1 (*n* = 3) were mixed with water at a ratio of 60 % w/v seaweed and 40% v/v of Milli-Q water in 120 mL sterile bottles. The minced slurry was then placed in an incubator at 37 °C. After the slurry reached a temperature of 30 °C (approximately 2 h), the seaweed solid fraction was separated from the liquid part by centrifugation at 64000 ×g for 30 min. The seaweed solid fraction was then rinsed with 500 mL of Milli-Q water and was again centrifuged under the same conditions. Samples were thereafter lyophilized, ground, and sieved (2 mm mesh) and stored at −20 °C until further analysis. The samples were made in triplicate, and the results were presented as mean ± standard deviation.

### 2.6 Analytical methods

#### 2.6.1 Bacterial growth and pH monitoring

Bacterial growth in selective media containing seaweed monosaccharides (Section 2.4) was measured by monitoring the change in optical density at 600 nm (OD_600_) using a spectrophotometer.

Additionally, the change in pH throughout the fermentations was measured using a pH meter.

#### 2.6.2 Total solid, moisture, and ash (mineral) content

Total solid, ash, and moisture content of the seaweed samples were determined according to Van Wychen S. and Laurens L.M.L. (19). In brief, a 2 g freeze-dried sample (*n* = 3) was added to preconditioned crucibles and placed in a heating oven (Fratelli Galli Control AG-System, Italy) set at 105 °C to reach a constant weight. Total solids were determined by weighing the samples. The same samples were thereafter combusted at 575 °C in a muffle furnace (Nabertherm Controller B180, Germany) for 3 h without ramp-up mode, and the ash (mineral) content was determined gravimetrically.

#### 2.6.3 Carbohydrate analysis

Carbohydrates, including neutral monosaccharides, mannitol, and uronic acids, were analyzed based on the method of Van Wychen and Laurens (20) for determining total carbohydrates in seaweed biomass from different treatments. In this method, 25 mg of freeze-dried seaweed powder (*n* = 3) was placed in glass tubes with screw lids and mixed with 250 µL of 72% (w/w) sulfuric acid. The samples were incubated at 30°C and 300 rpm for 60 minutes, then diluted to 4% (w/w) H_2_SO_4_ by adding Milli-Q water. Subsequently, the samples underwent heat treatment by autoclaving at 121°C for 60 minutes and centrifuging at 3,000 × g for 5 minutes to separate the supernatant from the solids. The final sample preparation step involved neutralizing with 0.1 M Ba(OH)_2_·H2O and filtering into analytical vials using 0.2 µm syringe filters.

The total carbohydrates in the seaweed samples and monosaccharides in MRS media were analyzed using a High-Performance Anion Exchange Chromatography system equipped with a Pulsed Amperometric Detector (HPAEC-PAD; Thermo Fisher Scientific, United States) and an analytical column and guard column (Dionex CarboPac PA-20; Thermo Fisher Scientific, Waltham, United States). Analyses were conducted under isocratic conditions with a flow rate of 0.5 mL/min at a temperature of 30°C. Monosaccharides and mannitol were eluted using mixtures of Milli-Q water (62.5%), 2 mM sodium hydroxide (37.5%), and 200 mM sodium hydroxide for 23 minutes. Uronic acids were eluted with mixtures of Milli-Q water (55%), 1 M sodium acetate in 200 mM sodium hydroxide (15%), and 200 mM sodium hydroxide (30%) for 15 minutes. Reference standards used included L(−)fucose, D (+) glucose, D (+)xylose, D (+) galactose, L(+) arabinose, D(+) mannose, D-mannitol, mannuronic acid, guluronic acid, and glucuronic acid.

#### 2.6.4 Quantification of organic acids

The content of lactic acid and short-chain fatty acids (SCFAs) in the liquid fraction from the lactic acid fermentation was analysed according to Allahgholi et.al. (21). The samples (*n* = 3) were centrifuged at 30000×g (Centrifuge 5425, Eppendorf) for 10 minutes, whereafter 1 mL of supernatant was mixed with 20 uL of 20 % (v/v) sulfuric acid and incubated for 30 minutes at 4 °C. Quantification was performed using the high-performance liquid chromatography (HPLC) system Dionex UltiMate 3000 RSLC (Thermo Fisher Scientific, US), equipped with an Aminex HPX-87H column (Bio-Rad, US).

#### 2.6.5 Protein analysis

Total protein content in seaweed samples was determined using the Dumas method by analysing the nitrogen content with a N/Protein Analyzer Flash EA 1112 Series (Thermo Electron Corp., Waltham, MA, USA) and Eager Smart software for data handling. Approximately 25 mg of freeze-dried sample (*n* = 3) was placed in aluminium containers for analysis. Aspartic acid was used as a reference for known protein content. Samples were combusted in a high-temperature chamber (800–1000 °C) with oxygen, producing carbon dioxide, water, and nitrogen. By-products were captured or separated in a column, and nitrogen content was determined by a thermal conductivity detector. A nitrogen-to-protein conversion factor of 5 was used to estimate protein content (22).

#### 2.6.6 Element analysis

Elemental content of seaweed samples (*n* = 3) was analysed by the accredited external laboratory ALS Scandinavia, as previously described by Jönsson et al. (2023)(23). Sample preparations were performed by heated wet digestion in nitric acid (HNO_3_) and hydrofluoric acid (HF) before quantification using high-resolution inductively coupled plasma mass spectrometry (HR-ICP-MS; Element XR, Thermo Scientific, Germany). Analytes detected from the method include arsenic (As), cadmium (Cd), calcium (Ca), chromium (Cr), cobalt (Co), cupper (Cu), iron (Fe), lead (Pb), magnesium (Mg), manganese (Mn), mercury (Hg), molybdenum (Mo), nickel (Ni), phosphorous (P), potassium (K), selenium (Se), silver (Ag), sodium (Na), and zinc (Zn),

The method differed slightly for the halogen iodine (I). Samples were prepared by microwave-assisted, closed vessel nitric acid (HNO_3_) digestion. The digests were thereafter diluted with a mixture of 0.5 % ammonia, 0.01 % Triton X-100, and 0.01 % EDTA before analysis by HR-ICP-MS.

#### 2.6.7 Antioxidant analysis

##### 2.6.7.1 Extraction of antioxidant compounds

Polyphenols, carotenoids, and tocopherols were extracted with a supercritical fluid extraction method optimised by Gondo et al 2023 (24). Briefly, 0.5 g of seaweed was weighed in a 5 mL extraction cell and subsequently filled with glass beads. The sample was extracted in three steps, employing the following conditions: Initially, CO_2_/EtOH/H_2_O (95/5/0, v/v/v) at 60°C, secondly, CO_2_/EtOH/H_2_O (35/60/5, v/v/v) at 70°C, and lastly, CO_2_/EtOH/H_2_O (15/72/13, v/v/v) at 70°C. This was done to comprehensively target nonpolar carotenoids and tocopherols and polar phenolic compounds. The extraction pressure was maintained at 300 bar, while the flow rate used was 3 mL/min. All extracts were then combined in a 100 mL bottle, followed by N_2_ drying to remove the organic solvent and freeze drying to remove the water. The dried extract was then reextracted with two solvent portions, i.e., 2 mL of water /methanol (70/30), then 2 mL of ethyl acetate/methanol (50/50). The first portion was investigated for phenolic acids and phloroglucinol, while the second portion was analysed for carotenoids and tocopherols. All extractions were done in triplicate (n=3).

##### 2.6.7.2 Quantification of antioxidant compounds

Analysis (*n* = 3) of phenolic acids, flavonoids, phloroglucinol, and tocopherols was performed by high-performance liquid chromatography with diode array and fluorescence detection (HPLC-DAD-FLD), using an Agilent HPLC 1100 series equipment, equipped with a Waters Xselect C-18 column (150 mm×3 mm, 3.5 µm). The following conditions were employed: DAD set at four different wavelengths being 260 nm for phloroglucinol, 280 nm for phenolic acids, 360 nm for flavonoids, and FLD set at 290 nm excitation and 330 nm emission for tocopherols; a mobile phase: (A) 1% aqueous formic acid (FA) and (B) 1% FA in acetonitrile, a column temperature of 25 °C, a flow rate of 0.5 mL/min, an injection volume of 20 µL; and a gradient: 0-5 min (2% B), 5-10 min (2-20% B), 10-15 min (20% B), 15-18 min (20-25% B), 18-20 min (25% B), 20-25 min (25-70% B), 25-30 min (70-100% B). The gradient was then kept at 100% B for up to 50 min to allow elution of tocopherols. β and γ-tocopherol were quantified as a mixture of isomers using the γ-tocopherol equivalence standard, as no separation was obtained for the two compounds.

The analysis method for carotenoids (*n* = 3) was adopted from Gondo et al. 2023 (24). Ultrahigh-performance supercritical fluid chromatography with diode array (UHPSFC-DAD) (Waters, Milford, MA, USA) equipped with a Waters Torus 1-aminoanthracene column (100 mm × 3 mm, 1.7 μm) was utilised under the following conditions; a mobile phase composition (A) CO_2_ and (B) methanol, with gradient composition as : 0-2.5 min (3% B), 2.5-5 min (12% B), 5-7 min (12-15% B), 7-9 min (15-25% B), 9-9.5 min (25% B), 9.5-10 min (25-3% B), 10-12 min (3% B). Column temperature was set at 30°C, a flow rate of 1.5 mL/min, an injection volume of 4 µL, and DAD set at 430 nm. The make-up solvent was 0.2% ammonium formate in methanol with a flow rate of 0.5 mL/min.

### 2.7 Statistics

Descriptive statistics were performed in Microsoft Excel (version 2510, WA, USA) and presented as mean value ± standard deviation (SD). To assess significant differences among treatments, one-way analysis of variance (ANOVA) and principal component analysis (PCA) were conducted using jamovi (version 2.6.44, Australia). Tukey’s *post hoc* test was utilized to identify significantly different sample groups. Statistics data can be found in Table S1-S8, including composition of untreated *A. esculenta* (Table S6) and the antioxidant composition of untreated and fermented *Alaria esculenta* (Table S7).

## 3 Results

### 3.1 Growth of *Lactiplanitbacillus plantarum* using mannitol or glucose as carbon sources

Single-substrate cultivations with mannitol or glucose in MRS medium were conducted to demonstrate the ability of the selected *L. plantarum* to convert relevant monosaccharides available in the brown seaweed biomass to lactic acid (Fig. 3A). The results showed that the strain utilized both mannitol and glucose (Fig. 3B), resulting in verified production of cells (monitored as an increase in optical density (OD)) and production of lactic acid. The OD_600_ at the end of the cultivation in fermenters containing mannitol and glucose were 9.2 and 14.5, respectively (Fig. 3C), showing a higher cell mass using glucose as carbon source, while the difference in pH due to lactic acid production (the major metabolite) was relatively small (Fig. 3D). Acetic acid was found in all cultivations (with mannitol, glucose, and when no carbon source was added), as a consequence of the MRS composition. However, the amount was lower in the mannitol-containing cultivation, compared to the cultivation in the presence of glucose and in the negative control (data not shown). This is in line with the results of Yang et al. 2019 (25), who reported that the use of sorbitol or mannitol as a carbon source resulted in acetate consumption involving aldehyde-alcohol dehydrogenase (AdhE). Mannitol has a lower redox potential (−0.32V) than glucose (−0.17V), and needs an extra oxidation step before entering glycolysis, accompanied by the formation of NADH (26). In the early stage, growth of *L. plantarum* on sorbitol or mannitol results in conversion to large amounts of lactic acid, with NADH accumulating in the cells, while in the later stage of growth, acetate is utilized to accelerate the consumption of NADH through AdhE (25). This redox imbalance, generated upon growth on mannitol, may be a reason for the lower amount of cell mass obtained at harvest.

**Figure 3.**
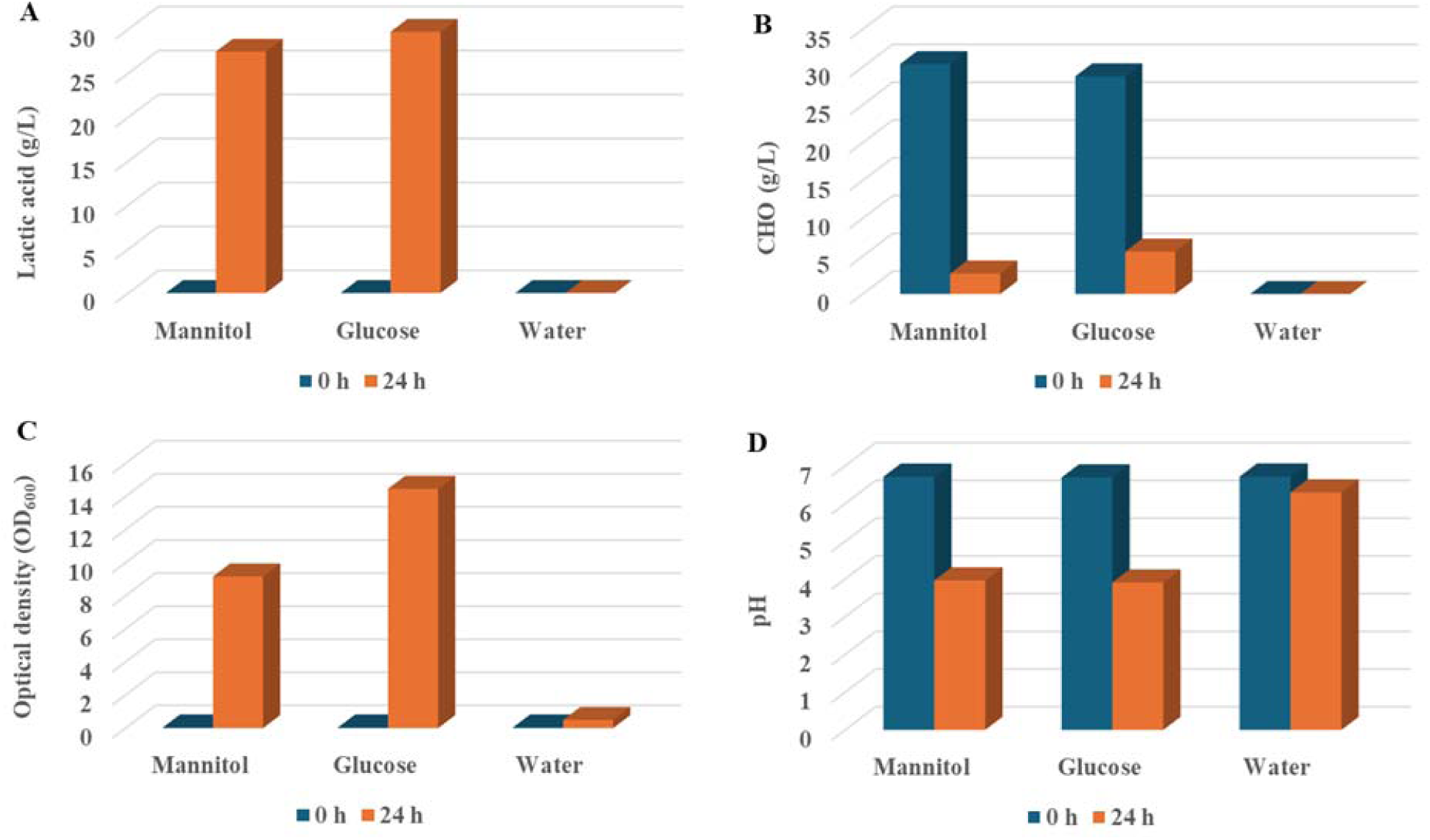
Summary of parameters for the single-substrate fermentation by *Lactiplantibacillus plantarum* in MRS. The lactic acid (A), carbohydrate content (CHO) (B), optical density (OD_600_) (C), and pH profile (D) at the start (0 h) and end of the cultivation (24 h) are shown.

### 3.2 Quality and food safety of *Alaria esculenta* after fermentation with *Lactiplantibacillus plantarum*

After confirmation of the utilization of mannitol, this main free sugar alcohol in *A. esculenta* (5.40 ±0.095 g/100 g DW), was targeted as a carbohydrate source in lactic acid fermentation of fresh froz n *A. esculenta*, using *L plantarum* as inoculum. This was performed in two modes of operation, submerged fermentation (SmF-LA) and solid state fermentation (SSF-LA), where LA stands for lactic acid which is expected as a main metabolite. Moreover, cultivation of any indigenous microbiota present on the seaweed batch was set up and run in parallel using the corresponding modes of operation (SmF-SP and SSF-SP, where SP stands for spontaneous fermentation by indigenous microbiota in the seaweed batch). The cultivations were made to investigate the effects of the respective type of fermentation on the seaweed’s nutritional quality and food safety. Nutritional quality is a measure of a well-balanced ratio of essential nutrients (carbohydrates, proteins, lipids, minerals, and vitamins) and was investigated by analyzing carbohydrate content, protein, organic acids, minerals, and antioxidant compounds. The lipid content in seaweed is low, motivating its exclusion from the current data. The food safety aspect was, apart from monitoring the decrease in pH, also targeted by analyzing potential toxic elements.

#### 3.2.1 Carbohydrate content and organic acid production

Lactic acid fermentation is a traditional preservative method for foods where the utilized carbohydrates are converted into organic acids, which leads to a lowering of the pH, limiting the growth of pathogens, so food can be stored for a longer period. To monitor changes in carbohydrate content, the total carbohydrate profile of the fresh frozen seaweed batch (untreated sample) was analyzed, and the analysis was again made on the same batch after fermentation (using the two modes of operation).

Subsequently, end-point samples from the two *L. plantarum* fermentation types (SmF-LA and SSF-LA) were collected, and the same was done for cultivations of indigenous microorganisms (SmF-SP and SSF-SP) that were run under the same conditions without inoculation, hence utilizing only indigenous microbiota from the samples.

Total carbohydrate content (Table 1, see also Fig.7A below) was highest in the untreated sample (16.6 ±1.4 g/100 g DW), which was significantly different compared to the carbohydrate content in fermented samples. Of the two inoculated samples, SmF-LA had the lowest total carbohydrate content at the end of the cultivation (10.51±0.44 g/100 g DW), while SSF-LA had a higher total carbohydrate content (after acid hydrolysis) at the end of the cultivation (14.26±0.55 g/100 g DW), indicating a lower efficiency in the utilization of available carbohydrates with this technique.

**Table 1.** Carbohydrate profile of *Alaria esculenta* with different treatments after acid hydrolysis. Results are reported as g/100 g ± standard deviation and are based on algal dry weight (DW). Significant differences (*p* < 0.05) among samples are indicated by letter (a-d). For statistical data, see Table S1.

| Carbohydrate (g/100 g DW) | Untreated | SmF-SP | SmF-LA | SSF-SP | SSF-LA | p-value |
| --- | --- | --- | --- | --- | --- | --- |
| Mannitol | 5.40±0.10<sup>a</sup> | 2.57±0.16<sup>b</sup> | 0.72±0.19<sup>d</sup> | 1.92±0.04<sup>c</sup> | 1.92±0.21<sup>c</sup> | < .001 |
| Fucose | 0.91±0.03<sup>b</sup> | 1.87±0.14<sup>a</sup> | 1.70±0.03<sup>a</sup> | 1.73±0.08<sup>a</sup> | 1.93±0.15<sup>a</sup> | < .001 |
| Glucose | 1.88±0.16<sup>c</sup> | 2.22±0.12<sup>bc</sup> | 3.79±0.10<sup>a</sup> | 2.69±0.71<sup>b</sup> | 3.11±0.16<sup>ab</sup> | < .001 |
| Xylose | 0.22±0.04<sup>b</sup> | 0.30±0.08<sup>ab</sup> | 0.25±0.01<sup>b</sup> | 0.32±0.05<sup>ab</sup> | 0.40±0.041<sup>a</sup> | 0.008 |
| Mannose | 0.03±0.03 | 0.02±0.01 | 0.02±0.01 | 0.02±0.00 | 0.02±0.004 | 0.898 |
| Glucuronic acid | 1.67±0.6107<sup>a</sup> | 0.98±0.23<sup>b</sup> | 0.67±0.04<sup>c</sup> | 0.86±0.46<sup>b</sup> | 1.77±0.29<sup>a</sup> | 0.016 |
| Guluronic acid | 1.03±0.07<sup>a</sup> | 0.63±0.09<sup>bc</sup> | 0.40±0.07<sup>c</sup> | 0.646±0.12<sup>b</sup> | 0.851±0.09<sup>ab</sup> | < .001 |
| Mannuronic acid | 5.44±0.97<sup>a</sup> | 4.33±0.45<sup>ab</sup> | 2.96±0.24<sup>b</sup> | 3.85±0.77<sup>ab</sup> | 4.26±0.19<sup>ab</sup> | 0.007 |
| Total CHO | 16.57±1.40<sup>a</sup> | 12.96±0.89<sup>bc</sup> | 10.51±0.44<sup>c</sup> | 12.03±1.8<sup>bc</sup> | 14.27±0.55<sup>ab</sup> | < .001 |

Mannitol was most efficiently used, and as expected, the mannitol content was highest in the untreated sample (5.40±0.095 g/100 g DW) and was significantly reduced after fermentation at all conditions (inoculated as well as conditions with indigenous microbiota). Statistically, the untreated sample was significantly different (p < 0.05) from all fermented samples, and the lowest carbohydrate content was recorded after SmF-LA (0.72±0.19 g/100 g DW). Utilization was less efficient using SmF-SP, SSF-SP, and SSF-LA as intermediate mannitol levels remained at the end of the fermentation.

Fucose content (originating from the polymer fucoidan) was, on the other hand, significantly higher in the fermented samples compared to the untreated one, indicating more efficient release after fermentation, resulting in more accessible material for total acid hydrolysis. Fucoidan also contains varying amounts of xylose, mannose, and glucuronic acid (27). The same trend as for fucose is seen for xylose, while glucuronic acid decreased, indicating that it may be hydrolyzed and consumed by the bacteria during the fermentation. Mannose content was low and did not show significant differences across treatments.

Glucose content (from the polymers laminarin and/or cellulose) also increased after fermentation, indicative of increased release by fermentation. Variation was, however, observed, with higher glucose content in *L. plantarum* cultivations than in cultivations of indigenous microbiota, indicating that either some hydrolysis of the polymer occurred in the latter case, followed by consumption, or that less of the polymer was released and remained less accessible.

Guluronic acid and mannuronic acid (the components of alginate) decreased somewhat after fermentation treatments, despite the lack of enzyme systems for alginate degradation in *L. plantarum* (based on genetic information of several sequenced strains, as indicated in Cazy [www.cazy.org])

Overall, the data show that fermentation significantly (*p* < 0.05) reduced the carbohydrates available for metabolism in *A. esculenta*, with a reduction in mannitol, while fucose and glucose content were concentrated across most treatments.

Both fermentation modes (SmF-LA and SSF-LA) reduced the pH of the *A. esculenta* seaweed slurries after inoculation with *L. plantarum*. According to Løvdal et al, (28) acidification down to pH 4.3 is sufficient for storage at refrigerated conditions, while a further decrease to pH 3.7 may be necessary for safe storage (preventing pathogenic microorganisms) at ambient temperatures. In this work, the desired pH for refrigerated storage was reached during the course of the fermentation between 48-72 h (Fig. 4A), dependent on the mode of operation. At 72h, the cultures were harvested. SmF-LA proved more effective than SSF-LA in decreasing the pH, which was reduced from 6.20 at inoculation to 4.04 in SmF-LA, while a reduction from 6.30 at inoculation to 4.25 was observed in SSF-LA. The difference may be due to a more efficient mixing of the cells with the seaweed substrate in SmF. The pH in the spontaneous fermentations (indigenous microorganisms from the seaweeds) did not change considerably (Fig. 4A), resulting in unsuitable conditions for preservation.

**Figure 4.**
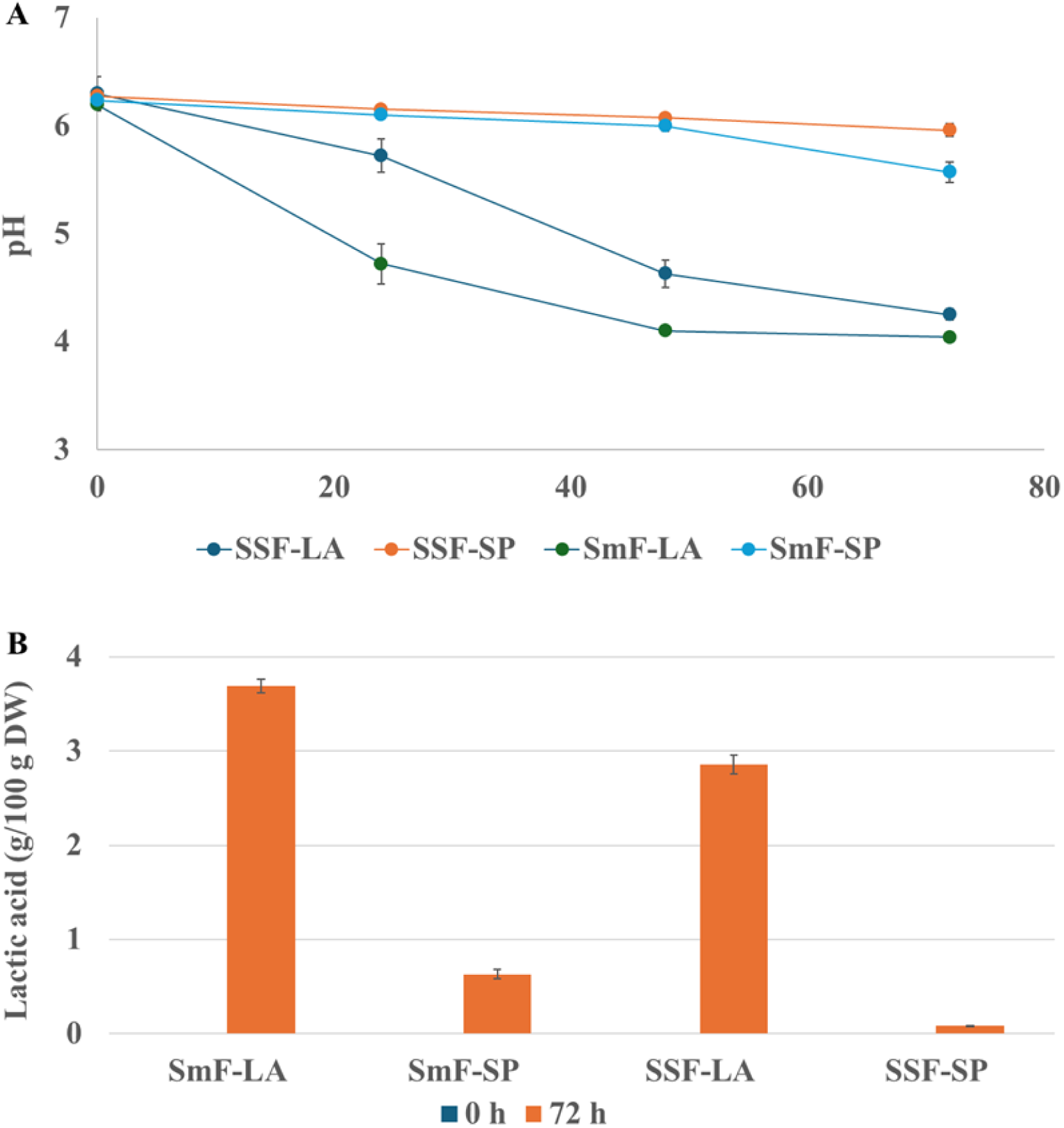
The pH profiles (A) and lactic acid production (B) during fermentation of brown seaweed with *Lactiplantibacillus plantarum* from different lactic acid fermentation systems, submerged fermentation (SmF-LA) and solid-state fermentation (SSF-LA), of the brown seaweed *Alaria esculenta,* compared to spontaneous fermentation by any indigenous microorganisms present in the seaweeds (SmF-SP and SSF-SP, respectively).

The organic acid content was verified by HPLC analysis at the start and end of the respective fermentation. At the end of the fermentation, lactic acid was the main organic acid in both SmF-LA and SSF-LA of *A. esculenta* (Fig. 4B), while only low amounts of lactic acid were detected in the SmF-SP and SSF-SP (Fig. 4B), showing that a minor production of organic acids occurred from indigenous microbiota, which, however, did not result in any major change of the pH in the culture. Acetic acid and formic acid were also produced in all fermentations, but at low levels (< 0.1 g/L, data not shown). Both fermentation modes inoculated with *L. plantarum* (SmF-LA and SSF-LA), thus resulted in successful lactic acid fermentation of *A. esculenta*, with sufficient decrease in pH for preservation at refrigerated conditions, motivating further analysis.

#### 3.2.2 Protein content

The protein content in the *A. esculenta* samples studied increased following the fermentations (Fig. 5, see also Fig.7A below). Untreated *A. esculenta* had a protein content of 11.6 ± 0.30 % DW, and fermentation under different conditions resulted in a significantly (*p* < .001) increased total protein content. While not at a significant level, the protein content after submerged fermentation with *L. plantarum* (SmF-LA 15.9±0.57 % DW) was highest, whereas the protein content in the indigenous microbiota cultivations (SmF-SP and SSF-SP with a protein content of 14.9±0.58 % DW and 14.9±0.65 % DW, respectively) and the SSF-LA (14.6±0.41% DW) were in the same range.

**Figure 5.**
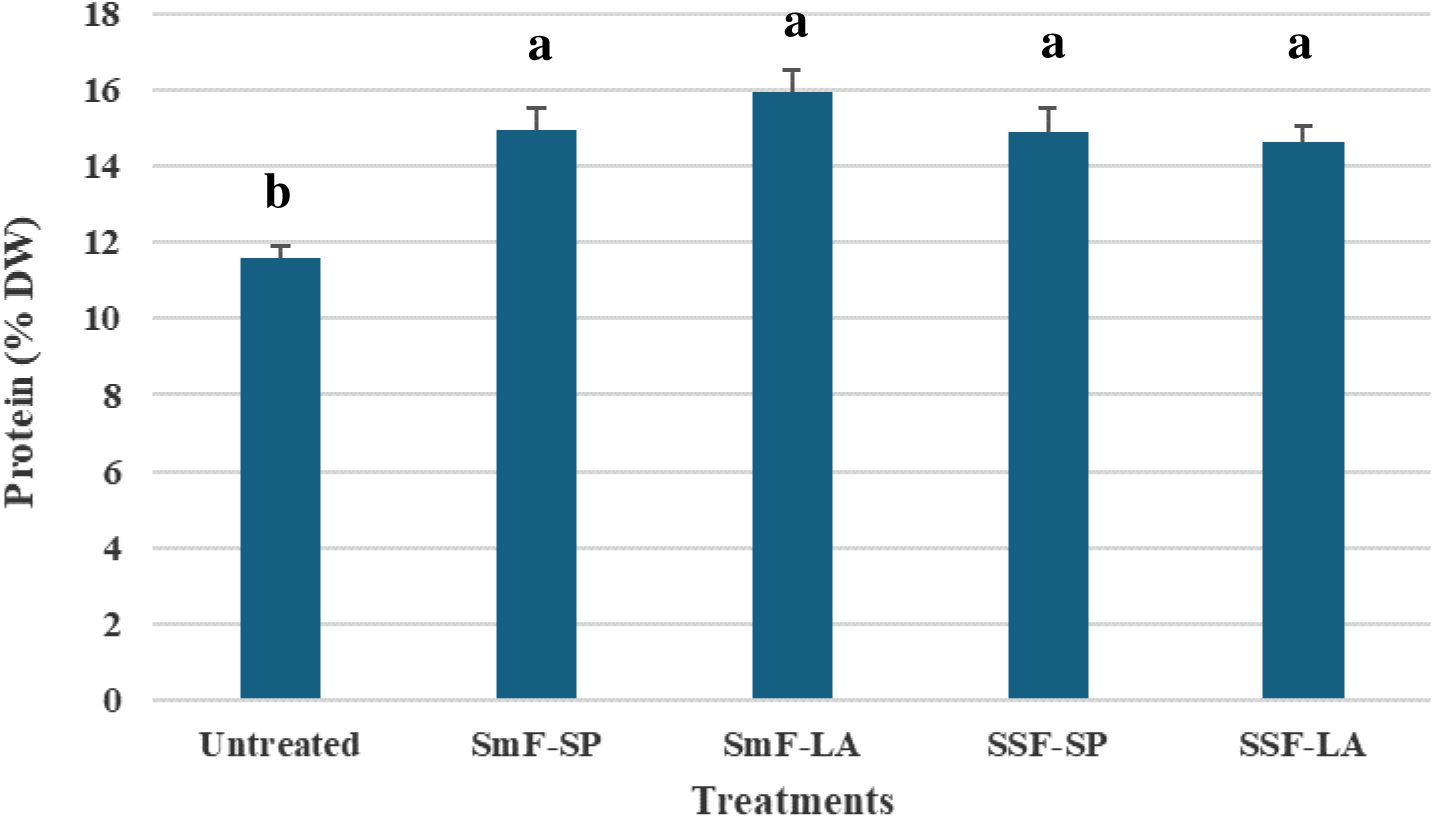
Percentage protein content of dry weight (*n* = 3) in untreated and fermented *Alaria esculenta* from SmF and SSF processes. Significant differences (Tukey’s HSD, *p* < 0.05) among samples are indicated by letter (a-b). For statistical data, see Table S2.

#### 3.2.3 Antioxidant compounds

The presence of various antioxidants was screened from the untreated *A. esculenta,* and *A. esculenta* fermented in the two SmF cultivations, i.e., *L. plantarum* (SmF-LA) and indigenous microbiota (SmF-SP). The major antioxidant compounds detected were phloroglucinol and fucoxanthin, which are typical compounds in brown seaweeds. Alterations in antioxidant compound concentrations as a consequence of the fermentation were clear, as shown by the trends in the heatmap (Fig. 6). For instance, there was a drastic increase in phloroglucinol content in the SmF-LA, compared to the untreated and to SmF-SP (Fig. 6), where the latter showed a less pronounced increase. This might indicate breakdown and release of more complicated polymeric phlorotannin structures into simple monomers by *L. plantarum*, but also to a certain extent by the indigenous strains, during fermentation. Larger polymers are generally difficult to extract, mostly because of solubility issues, but their conversion to small monomers could also enhance their extractability. Conversely, carotenoids showed a significant decrease after fermentation, indicating use by the microorganisms, while the α-tocopherol levels remained the same (Fig. 6) (*p* = 0.49).

**Figure 6.**
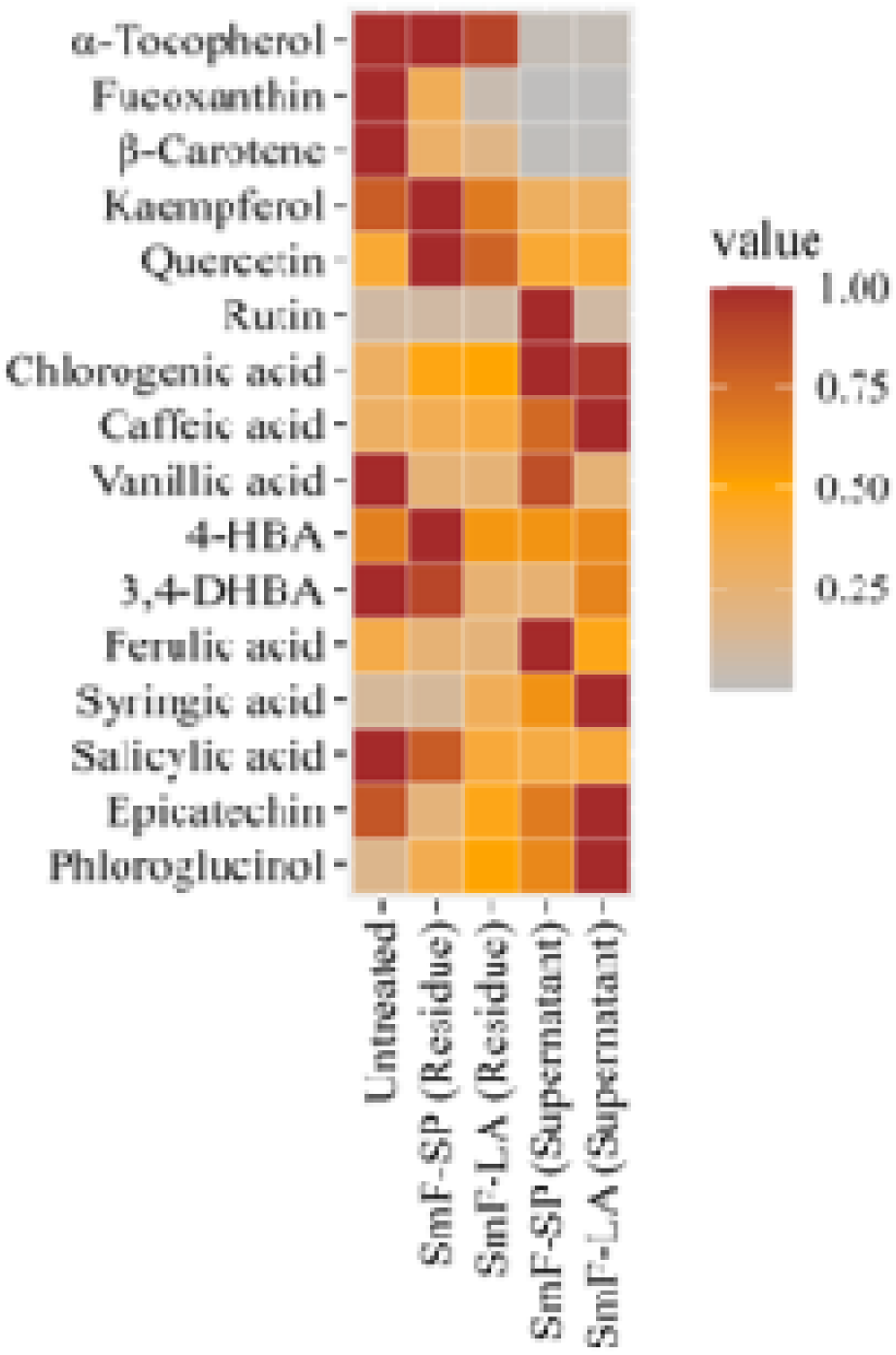
Heatmap plot showing the abundance of each antioxidant compound in the untreated seaweed, SmF-SP, and SmF-LA. The plot compares the compound levels in the solid residue and the supernatant obtained after submerged *Alaria esculenta* fermentation. Values in each row were normalized to their maximum concentration. For the original concentration data and statistical significance, see Table S7 and S8 in the Supplementary Material.

Although phenolic compounds were not analyzed in the SSF-LA and SSF-SP samples, they were screened in both the solid (residue) and the liquid fraction (supernatant) removed after fermentation in the SmF-SP and SmF-LA samples. This data provided insights into how water influences the leaching of antioxidant compounds to the aqueous phase in the SmF samples, a phenomenon unlikely to occur to the same extent in the SSF samples. As expected, a clear trend emerged, where water-soluble compounds such as phenolic acids and phloroglucinol were present in high amounts in the supernatant of the submerged *A. esculenta* samples (i.e., both SmF-SP and SmF-LA). This is likely due to leaching into the liquid phase during fermentation. Conversely, for SmF-SP, 3,4-DHBA and kaempferol were only detected in the solid residue, while their concentrations were below the limits of quantification in the liquid fraction. Although these compounds generally have good water solubility, some may remain more strongly bound to the matrix, affecting their leaching kinetics.

Non-polar compounds such as tocopherols and carotenoids were not detected in the liquid fraction, as expected (Fig. 6). In general, the concentrations of many phenolic compounds were lower than those of phloroglucinol and carotenoids (Table S7). Notably, the concentrations of phenolic acids, including ferulic acid, syringic acid, caffeic acid, and chlorogenic acid, increased in SmF-LA and SmF-SP samples (compared to untreated *A. esculenta* samples), when the amounts in both the residue and supernatant were considered. This suggests either metabolic transformation or enhanced extractability post-fermentation. Additionally, epicatechin and syringic acid levels increased in the SmF-LA samples compared to both the untreated *A. esculenta* samples and SmF-SP samples (Fig. 6).

#### 3.2.4 Mineral content

The effect of SmF-LA and SSF-LA on mineral content in the solid *A. esculenta* fraction was evaluated and compared to fresh frozen seaweed (untreated) and water-soaked control samples. Both soaking and fermentation reduced the total mineral (ash) content (37.56 ± 0.60 % of DW) significantly (*p* < 0.05), with an approximately two-fold reduction (Table 2). The total mineral fraction of the seaweed samples was dominated by five elements: K, Na, Ca, Mg, and P (Table 3, Fig. 7B)). Whereas K, Na, and Mg were significantly reduced after all treatments, Ca was concentrated, and P remained unaffected.

**Figure 7.**
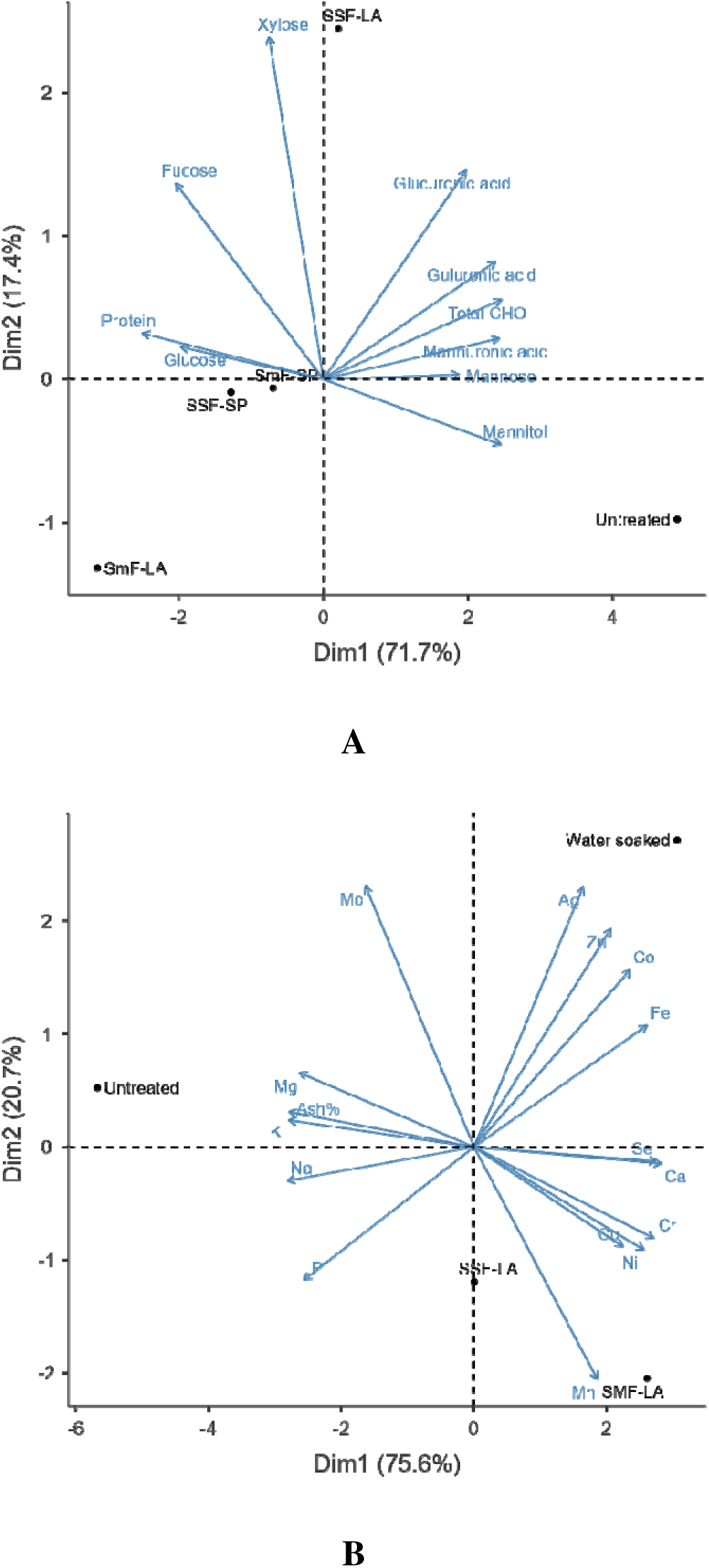
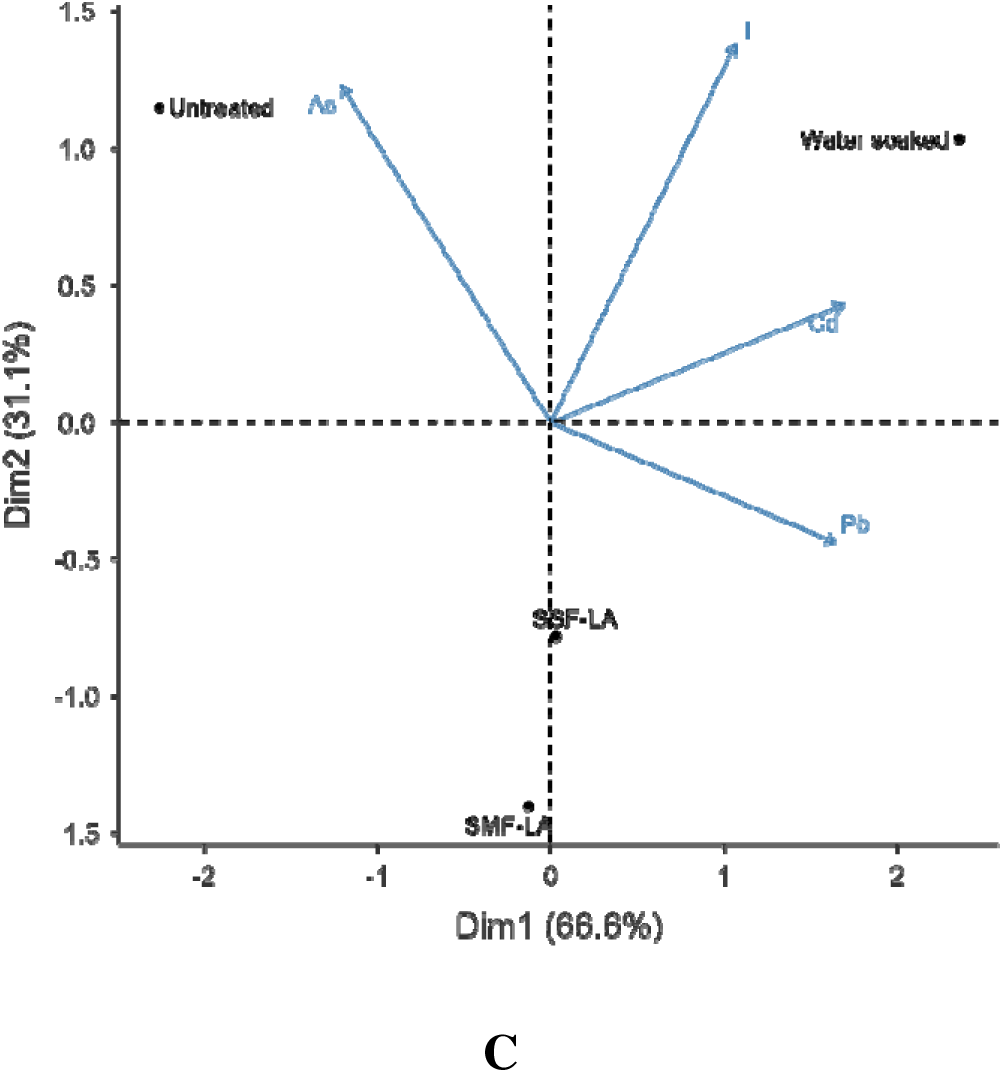
Principal component analysis (PCA) biplots illustrating the impact of A) monosaccharides and protein, B) minerals, and C) potentially toxic elements on sample similarities.

**Table 2.** Total solids and ash (mineral) content of the seaweed sample after water soaking or fermentation. The data (*n* = 3) is presented as mean values with standard deviation, and significant differences (Tukey’s HSD, *p* < 0.05) between samples are indicated by letters (a–c). For statistical data, see Table S3.

| Treatments<br>(g/ 100 g DW) | Untreated | Water soaked | SmF-LA | SSF-LA | *p*-value |
| --- | --- | --- | --- | --- | --- |
| **Total solids** | 91.58 ± 0.09 | 93.34 ± 2.69 | 90.74 ± 4.13 | 92.98 ± 1.45 | 0.593 |
| **Total minerals (ash)** | 37.57 ± 0.60 <sup>a</sup> | 15.96 ± 1.39 <sup>c</sup> | 15.84 ± 1.16 <sup>c</sup> | 19.64 ± 1.84 <sup>b</sup> | <.001 |

**Table 3.** Mineral content in *Alaria esculenta* subjected to water soaking or fermentation. The data (*n* = 3) is presented as mean values with standard deviation, and significant differences (Tukey’s HSD, *p* < 0.05) between samples are indicated by letters (a–d). For statistical data, see Table S4.

| Elements<br>(mg/kg DW) | Untreated | Water soaked | SmF-LA | SSF-LA | *p*-value |
| --- | --- | --- | --- | --- | --- |
| K | 79 300 ± 2 443 <sup>a</sup> | 42 233 ± 2 503 <sup>bc</sup> | 38 400 ± 3 503 <sup>c</sup> | 56 900 ± 11 811 <sup>b</sup> | <.001 |
| Na | 48 167 ± 1 966 <sup>a</sup> | 13 902 ± 10 662 <sup>b</sup> | 18 700 ± 2 206 <sup>b</sup> | 29 200 ± 6 061 <sup>b</sup> | <.001 |
| Ca | 11 533 ± 723 <sup>b</sup> | 14 567 ± 1 050 <sup>a</sup> | 14 600 ± 346 <sup>a</sup> | 13 567 ± 850 <sup>ab</sup> | 0.004 |
| Mg | 11 567 ± 493 <sup>a</sup> | 7 705 ± 602 <sup>b</sup> | 7 393 ± 385 <sup>b</sup> | 7 440 ± 79 <sup>b</sup> | <.001 |
| P | 3 817 ± 146 | 3 269 ± 243 | 3 490 ± 265 | 3 640 ± 230 | 0.084 |
| Fe | 127 ± 6 <sup>c</sup> | 205 ± 12 <sup>a</sup> | 172 ± 11 <sup>b</sup> | 164 ± 6 <sup>b</sup> | <.001 |
| Zn | 51 ± 2 <sup>b</sup> | 86 ± 5 <sup>a</sup> | 61 ± 6 <sup>b</sup> | 56 ± 4 <sup>b</sup> | <.001 |
| Mn | 6.90 ± 0.09 <sup>c</sup> | 7.69 ± 0.66 <sup>bc</sup> | 10.05 ± 0.65 <sup>a</sup> | 8.45 ± 0.67 <sup>b</sup> | <.001 |
| Cu | 3.29 ± 0.48 <sup>b</sup> | 5.60 ± 0.33 <sup>a</sup> | 5.78 ± 0.38 <sup>a</sup> | 6.60 ± 0.32 <sup>a</sup> | <.001 |
| Ni | 1.75 ± 0.06 <sup>b</sup> | 1.99 ± 0.08 <sup>ab</sup> | 2.13 ± 0.16 <sup>a</sup> | 1.91 ± 0.09 <sup>a</sup> | 0.012 |
| Cr | 1.36 ± 0.18 | 1.64 ± 0.07 | 1.72 ± 0.22 | 1.62 ± 0.18 | 0.130 |
| Co | 0.123 ± 0.005 <sup>c</sup> | 0.201 ± 0.009 <sup>a</sup> | 0.158 ± 0.015 <sup>b</sup> | 0.146 ± 0.005 <sup>bc</sup> | <.001 |
| Se | 0.306 ± 0.015 <sup>b</sup> | 0.417 ± 0.023 <sup>a</sup> | 0.429 ± 0.046 <sup>a</sup> | 0.361 ± 0.044 <sup>ab</sup> | 0.009 |
| Mo | 0.205 ± 0.004 <sup>a</sup> | 0.194 ± 0.012 <sup>a</sup> | 0.143 ± 0.006 <sup>b</sup> | 0.165 ± 0.031 <sup>ab</sup> | 0.008 |
| Ag | 0.020 ± 0.002 | 0.185 ± 0.231 | 0.034 ± 0.001 | 0.039 ± 0.003 | 0.328 |

#### 3.2.5 Potentially toxic elements (PTEs)

The chemical safety of seaweed exerts a critical factor for its introduction and establishment into new food markets. Hence, potentially toxic elements (Table 4, see also Fig.7C below), including As, Cd, Hg, Pb, and I, should be monitored and measured to meet the limits, which today are mainly based on acceptance criteria by ANSES and the European Commission (16), and tolerable intake levels by EFSA. While iodine is an essential mineral in the right dose, excessive intake can pose negative health effects. In this study, concentrations of PTEs in fresh *A. esculenta* were analyzed in the solid fraction and compared with water-soaked and fermented samples (SmF-LA and SSF-LA). The amounts of Hg, Pb, and I in all samples were lower than the acceptance limitations. Only Cd exceeded the proposed acceptance limit. Total As does not currently have any acceptance limits, but was reduced by both soaking and fermentation. SmF-LA resulted in the largest reduction of the levels of I and As, whereas the amounts of Cd and Pb did not change significantly by fermentation. Interestingly, the results after water soaking differed from the fermentation samples by introducing a concentrating effect on I and Cd. Hg was found to be below the quantification limit (0.020 mg/kg DW) for all samples.

**Table 4.** Summary of potential toxic elements in *Alaria esculenta* subjected to different treatments. The data (*n* = 3) is presented as mean values with standard deviation, and significant differences (Tukey’s HSD) between samples are indicated by letters (a–c). For statistical data, see Table S5.

| Potentially toxic elements (mg/kg DW) | Untreated | Water soaked | SmF-LA | SSF-LA | *p*-value | Limit * |
| --- | --- | --- | --- | --- | --- | --- |
| I | 808 ± 35<sup>b</sup> | 1 380 ± 15<sup>a</sup> | 480 ± 35<sup>c</sup> | 594 ± 129<sup>c</sup> | <.001 | 2 000 |
| As | 50.7 ± 0.5<sup>a</sup> | 39.2 ± 4.7<sup>ab</sup> | 35.5 ± 4.7<sup>b</sup> | 40.1 ± 5.6<sup>ab</sup> | 0.014 | N/A (iAs:3) |
| Cd | 1.37 ± 0.06<sup>b</sup> | 2.20 ± 0.14<sup>a</sup> | 1.60 ± 0.09<sup>b</sup> | 1.62 ± 0.17<sup>b</sup> | <.001 | 0.5 |
| Pb | 0.156 ± 0.003<sup>b</sup> | 0.280 ± 0.055<sup>a</sup> | 0.229±0.010<sup>ab</sup> | 0.257 ± 0.025<sup>a</sup> | 0.005 | 5 |
| Hg | < 0.020 | < 0.020 | < 0.020 | < 0.020 | N/A | 0.1 |

\* ANSES (16)

## 4 Discussion

Fermentation using lactic acid bacteria is a promising bio-preservation technique for underutilized marine resources for food and nutraceutical applications. This study has demonstrated the potential of lactic acid fermentation in improving the quality and food safety of the brown algae, *A. esculenta,* for food applications.

Initially, the fermentability of a single-strain commercial LAB of the species *L. plantarum* on MRS media containing brown algae monosaccharides, mannitol, and glucose, was verified. Although mannitol utilization resulted in lower biomass production than when using glucose as a carbohydrat source, the lactic acid production and subsequently the desired pH reduction were comparable. Moreover, as explained in results, the lower biomass production is in line with literature data (21,25,26) as mannitol has a lower redox potential (−0.32V) than glucose (−0.17V), and needs an extra oxidation step before entering the glycolysis, resulting in a redox imbalance with NADH accumulating in the cells requiring acetate utilization via AdhE to accelerate consumption of NADH.

In this study, quantification of biomass production and kinetic growth parameters during seaweed fermentation was not possible due to the heterogeneous and viscous nature of the fermented seaweed slurry, which interfered with optical density measurements. This limitation has been reported in other complex fermentation systems where OD readings are unreliable (29). Future work should employ alternative approaches such as dry cell weight determination, automated OD assays, or soft sensor technologies to enable accurate monitoring of microbial growth and substrate consumption in seaweed-based fermentations.

Based on the verified ability of the *L. plantarum* culture to proliferate in mannitol supplemented MRS medium, fermentation using seaweed biomass was investigated using two modes of operation, submerged and solid-state fermentation (SmF-LA and SSF-LA, respectively), and compared with parallel cultivations without added bacterial culture (spontaneous fermentation; SmF-SP and SSF-SP, respectively, with growth of indigenous microbiota from the seaweed batch). Both SmF-LA and SSF-LA resulted in the desired pH reduction, below pH 4.3, reported to be crucial for food safety in refrigerated products (28), while this was not achieved when the indigenous microbiota proliferated. Generally, a seaweed product can be considered safe for consumption if the pH is below 4.3 if stored at temperatures up to 4 °C (cf. *Bacillus cereus*), or below 3.7 if stored at ambient temperatures (cf. *Salmonella* spp.) (28). Hence, the present study demonstrates successful lactic acid fermentation of *A. esculenta* by the *L. plantarum* bacterial culture using both submerged and solid-state fermentation, which indicates that these fermentation systems can serve as useful preservation methods of brown seaweed species (with mannitol as carbohydrate source, as discussed below) to control their microbial quality. Though this environment provides suitable preservative conditions for seaweed, other components in the biomass may be changed by microbial activity.

The carbohydrate profile of *A. esculenta* after different lactic acid fermentation treatments showed significant alterations, highlighting the impact of fermentation on the biochemical composition. Mannitol, a sugar alcohol that serves as a key storage carbohydrate in brown algae, was highest in the untreated sample, aligning with previous studies, showing its abundance in brown seaweeds like *Alaria esculenta* (30). All fermentation treatments resulted in a significant reduction in mannitol content, with the lowest levels observed after SmF-LA, confirming consumption during fermentation. LAB are known to metabolize mannitol under anaerobic conditions, explaining the marked decrease (21). Fucose, a key monosaccharide in fucoidans, was notably higher in the fermented samples, particularly in SSF-LA. The increase in fucose content suggests that lactic acid fermentation enhances the release of fucoidan from the seaweed structures, possibly due to enzymatic activities of the LAB or via breakdown of substituents or interactions in or between complex polysaccharides (31). Xylose showed moderate increases after fermentation, with the highest levels observed in the SSF-LA treatment. Xylose is a component in fucoidan polysaccharides in brown seaweeds, and its increase subsequently related to the increase in fucose. In line with the increased fucose content, glucose content also increased significantly, indicating a more efficient release from the seaweed material as a consequence of the fermentation. However, in this case, a certain hydrolysis of glucans may also occur during the fermentation, degrading glucose-containing polysaccharides like laminarin, into simpler glucose units (32–34), which may be consumed by the bacteria. This seems to be a possibility for the indigenous microbiota cultivations (with currently undefined enzyme systems) that show lower levels of glucose in the total hydrolysis than the *L. plantarum* cultivations (although higher than the untreated control). The guluronic acid and mannuronic acids, which are components of alginate, a major structural polysaccharide in brown algae (35), also showed variability after fermentation. The reduction in these acids in most fermentation treatments could suggest partial degradation or transformation of alginate, possibly due to the acidic environment created by the lactic acid bacteria (36). Overall, the total carbohydrate content was reduced in all fermentation treatments compared to the untreated sample, with the SmF-LA treatment showing the lowest carbohydrate levels. This aligns with previous findings that lactic acid fermentation can lead to the consumption and conversion of carbohydrates, especially mannitol, by LAB, resulting in lower overall carbohydrate content (4, 21, 37). However, the relatively higher carbohydrate content in the SSF-LA treatment suggests that solid-state fermentation is less effective on seaweed structures, resulting in less pronounced growth as compared to SmF.

Seaweed can be relatively rich in protein (38, 39) and 10-15 % of protein is normally found in dry brown seaweeds. This work revealed that the fermentation increased the overall level due to the addition of bacterial protein. The result showed that a higher percentage of protein was found in the fermented samples (both those inoculated with *L. plantarum* and those fermented by indigenous microbiota only) when compared to the fresh frozen untreated seaweed samples.

Overall, the changes in antioxidant compounds are complex and diverse. Most compounds exhibited similar trends between the control samples, where indigenous microbiota proliferated, and those fermented with *L. plantarum*. This highlights the role of native microbiota in influencing phenolic content. However, certain compounds, such as phloroglucinol, epicatechin, and syringic acid, were more prominent in the *L. plantarum*-fermented samples compared to the control, suggesting that this type of LAB plays a significant role in enhancing these compounds. Conversely, salicylic acid was more abundant in the control samples than in both the LAB-treated samples and the untreated seaweed. However, it remains to be determined whether the observed changes result from enzymatic transformations or enhanced extractability due to matrix changes. However, improved radical scavenging ability has been reported after fermentation of several seaweed species(40), mostly associated with increased phenolic compound content. Metabolic conversion of phenolic compounds during LAB fermentation by the action of esterase, decarboxylation and reduction has been reported by Gaur & Gänzle (41). These mechanisms could also suggest hydrolysis of large and polymeric phenolic compounds into small free phenolic compounds, which are easily extractable due to their high solubility in water. Sanchez-Maldonaldo et al (42) suggested conversion of hydroxycinnamic acids by LAB could be associated with an action to reduce the antibacterial activity of the phenolic compounds. It is also suggested that hydroxycinnamic acids, such as syringic acid, exhibit more antimicrobial activity compared to hydroxybenzoic acids. Other compounds may be decreased due to bacterial consumption during fermentation. For instance, vanillic acid was only observed with the non-fermented samples, while none was present in the non-induced control (only allowing indigenous microorganisms to grow) and the *L. plantarum* fermented seaweed. Interestingly, all the lipophilic compounds indicated a decrease in concentration after fermentation (both indigenous and *L. plantarum* fermented seaweed). Some studies suggest that lipophilic compounds are soluble in the bacterial membranes (42, 43), which therefore increases their consumption. However, carotenoids are also generally sensitive to light, therefore, the influence of such factors during the fermentation processes cannot be ruled out. On the other hand, an increase in carotenoid concentrations after fermentation has been reported in some studies, suggesting the influence of enzymatic activity in extractability (44, 45). In general, the diverse changes in antioxidant compound compositions as a result of fermentation might be dependent on the starting material and strain used.

Seaweed contains high levels and a wide diversity of minerals due to their marine habitat. Important minerals, such as calcium, accumulate in seaweed at much higher levels than in terrestrial food sources. In this study, the ash content of fresh seaweed (37.57 %) was drastically reduced after all treatment methods, which indicates that a large fraction of the elements associates weakly with the seaweed biomass. This agrees with previous studies by Jönsson & Nordberg Karlsson (23). The ash concentration after SmF-LA (15.84 %) was similar to that of water-soaked samples (15.96 %), while SSF-LA had slightly higher content (19.64 %). Both untreated and treated *A. esculenta* proved to be rich in potassium, sodium, calcium, magnesium, and phosphorus. However, to meet the intake requirements according to the Nordic Nutrition Recommendations (46), relatively large portions (25-160 g) of dry seaweed should be consumed daily. Solid state fermentation (SSF-LA) proved better at retaining the content of potassium and silver (while at low levels), whereas submerged fermentation (SmF-LA) kept higher manganese levels. There was no significant difference between the two fermentation systems for the remaining elements.

Among the essential elements, *A. esculenta* can be a good source of iodine. However, while proven to be a good source, only 0.1-0.2 g is required to satisfy the daily adequate intake level, and care should be taken not to exceed the tolerable upper level of 0.4-1.0 g per day. Similarly, the cadmium concentration should be considered when introducing seaweed as a food source in European markets. Up to 11-18 g of the studied seaweed can safely be consumed according to EFSA, although cadmium levels in all seaweed samples exceeded the acceptance levels by ANSES (16). Based on these results and current regulations, iodine and cadmium are the main concerns for seaweed as a food source in Europe, which agrees with previous studies (23, 47). Worth considering is that the alginate polysaccharide binds certain metal ions very well, which affects the bio-absorption of heavy metals through the human system and has been linked to the reduction of mineral availability due to the binding nature of the polysaccharide (22).

Previous studies have indicated that fermentation can become useful for the reduction of PTEs. For instance, Bruhn et al. (15) reported that LAB fermentation reduced the levels of cadmium, total arsenic, and mercury by 35.3, 5.9, and 34.8 %, respectively. Our study demonstrated a similar reduction in total arsenic (−21.0 to −30.0 %), but, contrary to Bruhn et al. (2023), showed an increase in cadmium (17.0 to 60.6 %) and lead (46.9 to 80.1 %) concentrations. Interestingly, while fermentation of seaweed reduced its iodine levels by 26.5 % using solid-state and 40.6 % by submerged fermentation, water-soaking instead increased the iodine content. Previous studies on brown seaweed show varying results on iodine content after soaking, with both increased content (23, 48) and reduced levels (49, 50). Overall, the two fermentation systems exerted a similar impact on the seaweed’s element content.

While this study demonstrates the improved nutritional quality of fermented seaweed, it did not comprehensively evaluate sensory attributes (taste, flavor, odor, appearance, and texture), shelf life, or probiotic viability post-fermentation. These parameters are critical for validating food application claims and should be considered in future investigations. Supporting evidence from a recent sensory study (51) shows that fermentation of *Alaria esculenta* with lactic acid bacteria reduces undesirable “seafood-like” flavors and enhances the perceived product freshness and consumer acceptance, underscoring the importance of incorporating such evaluations in subsequent work. These findings were reinforced by a recent review (52), which showed that bacterial fermentation mitigates off-flavors such as bitterness and marine notes, while enhancing pleasant aromas and improving overall sensory appeal.

All above results underline the profound impact of lactic acid fermentation on the composition of carbohydrates, organic acids, protein content, antioxidant compounds, minerals and PTEs of *A. esculenta*, with potential implications for its nutritional and functional properties. Consequently, lactic acid fermentation could be a valuable tool in optimizing the health-promoting properties of seaweeds for use in functional foods or nutraceuticals.

## 5. Conclusion

The selected *L. plantarum* showed the ability to utilize mannitol, the main sugar alcohol in *A. esculenta,* and glucose to produce cells and lactic acid. Lactic acid fermentation by the selected *L. plantarum* on fresh frozen *A. esculenta* affected the total amount and composition of carbohydrates in seaweed when compared to untreated samples. The data indicate that lactic acid fermentation significantly affected the carbohydrate profile of *A. esculenta*, with reductions in mannitol and increases in fucose and glucose content across most treatments. Fermentation with the addition of *L. plantarum* (SSF-LA and SmF-LA) increased the lactic acid concentration, resulting in reduced pH of the seaweed slurry. Fermentation with *L. plantarum* also increased the total protein content and showed no negative effect on the seaweed protein level. Fermentation also altered the concentrations of different antioxidant compounds. Lactic acid fermentation (SmF-LA) significantly increased the concentration of phenolic compounds, such as phloroglucinol, syringic acid, and epicatechin, compared to the untreated sample. However, lipophilic compounds, like carotenoids, decreased following both spontaneous- and lactic acid fermentation (SmF-SP and SmF-LA). The main limitation of this study is the lack of data on the changes in antioxidant composition within the solid-state fermentation matrix. However, the influence of water on the leaching of antioxidant compounds was clarified through analysis of the supernatant in the SmF samples. Future research should investigate whether these changes in antioxidant compounds composition are due to metabolic transformations or structural modifications that enhance extractability. Lactic acid fermentation of *A. esculenta* (SmF-LA and SSF-LA) as well as water soaking treatment significantly reduced the total mineral content, with key minerals like K, Na, and Mg decreasing, while calcium increased, and phosphorus concentrations remained unchanged. Submerged fermentation with the addition of *L. plantarum* (SmF-LA), effectively improved the removal of arsenic and iodine levels, though cadmium still exceeded acceptable limits. Soaking seaweed in water unexpectedly increased cadmium and iodine concentrations. Overall, fermentation using lactic acid bacteria showed potential as a bio-preservation method for the edible brown seaweed *A. esculenta,* compared to spontaneous fermentation, and especially SmF-LA is a promising fermentation process that can improve the nutritional profile and increase seaweed safety by lowering potentially toxic elements.

## Acknowledgments

The authors would like to thank Frans-Peder Nilsson and Emma Mathies for assistance in receiving and sending materials for analysis.

