## supplemental for "Fermentation of the Edible Brown Seaweed *Alaria esculenta* by *Lactiplantibacillus plantarum* affects nutritional profile and the content of potentially toxic elements"

**Table S1. Statistics for carbohydrate analysis (*n* = 3).**

| One-Way ANOVA (Fisher's) | | | | |
| --- | --- | --- | --- | --- |
|  | **F** | **df1** | **df2** | **p** |
| **Mannitol** | 394.110 | 4 | 10 | <.001 |
| **Fucose** | 52.047 | 4 | 10 | <.001 |
| **Glucose** | 14.779 | 4 | 10 | <.001 |
| **Xylose** | 6.340 | 4 | 10 | 0.008 |
| **Mannose** | 0.258 | 4 | 10 | 0.898 |
| **Glucuronic acid** | 5.171 | 4 | 10 | 0.016 |
| **Guluronic acid** | 23.101 | 4 | 10 | <.001 |
| **Mannuronic acid** | 6.649 | 4 | 10 | 0.007 |
| **Total Carb** | 12.154 | 4 | 10 | <.001 |

**Post Hoc Tests**

| Tukey Post-Hoc Test – Mannitol | | | | | | |
| --- | --- | --- | --- | --- | --- | --- |
|  |  | **SMF-SP** | **SSF-SP** | **SMF-LA** | **SSF-LA** | **Untreated** |
| **SMF-SP** | **p-value** | — | 0.003 | <.001 | 0.003 | <.001 |
| **SSF-SP** | **p-value** |  | — | <.001 | 1.000 | <.001 |
| **SMF-LA** | **p-value** |  |  | — | <.001 | <.001 |
| **SSF-LA** | **p-value** |  |  |  | — | <.001 |
| **Untreated** | **p-value** |  |  |  |  | — |

| Tukey Post-Hoc Test – Fucose | | | | | | |
| --- | --- | --- | --- | --- | --- | --- |
|  |  | **SMF-SP** | **SSF-SP** | **SMF-LA** | **SSF-LA** | **Untreated** |
| **SMF-SP** | **p-value** | — | 0.477 | 0.293 | 0.952 | <.001 |
| **SSF-SP** | **p-value** |  | — | 0.994 | 0.193 | <.001 |
| **SMF-LA** | **p-value** |  |  | — | 0.107 | <.001 |
| **SSF-LA** | **p-value** |  |  |  | — | <.001 |
| **Untreated** | **p-value** |  |  |  |  | — |

| Tukey Post-Hoc Test – Glucose | | | | | | |
| --- | --- | --- | --- | --- | --- | --- |
|  |  | **SMF-SP** | **SSF-SP** | **SMF-LA** | **SSF-LA** | **Untreated** |
| **SMF-SP** | **p-value** | — | 0.496 | 0.001 | 0.058 | 0.721 |
| **SSF-SP** | **p-value** |  | — | 0.016 | 0.562 | 0.089 |
| **SMF-LA** | **p-value** |  |  | — | 0.173 | <.001 |
| **SSF-LA** | **p-value** |  |  |  | — | 0.008 |
| **Untreated** | **p-value** |  |  |  |  | — |

| Tukey Post-Hoc Test – Xylose | | | | | | |
| --- | --- | --- | --- | --- | --- | --- |
|  |  | **SMF-SP** | **SSF-SP** | **SMF-LA** | **SSF-LA** | **Untreated** |
| **SMF-SP** | **p-value** | — | 0.980 | 0.739 | 0.145 | 0.315 |
| **SSF-SP** | **p-value** |  | — | 0.438 | 0.313 | 0.146 |
| **SMF-LA** | **p-value** |  |  | — | 0.022 | 0.920 |
| **SSF-LA** | **p-value** |  |  |  | — | 0.006 |
| **Untreated** | **p-value** |  |  |  |  | — |

| Tukey Post-Hoc Test – Mannose | | | | | | |
| --- | --- | --- | --- | --- | --- | --- |
|  |  | **SMF-SP** | **SSF-SP** | **SMF-LA** | **SSF-LA** | **Untreated** |
| **SMF-SP** | **p-value** | — | 1.000 | 0.995 | 0.976 | 0.893 |
| **SSF-SP** | **p-value** |  | — | 0.999 | 0.990 | 0.931 |
| **SMF-LA** | **p-value** |  |  | — | 1.000 | 0.983 |
| **SSF-LA** | **p-value** |  |  |  | — | 0.997 |
| **Untreated** | **p-value** |  |  |  |  | — |

| Tukey Post-Hoc Test – Glucuronic acid | | | | | | |
| --- | --- | --- | --- | --- | --- | --- |
|  |  | **SMF-SP** | **SSF-SP** | **SMF-LA** | **SSF-LA** | **Untreated** |
| **SMF-SP** | **p-value** | — | 0.994 | 0.848 | 0.153 | 0.250 |
| **SSF-SP** | **p-value** |  | — | 0.971 | 0.083 | 0.141 |
| **SMF-LA** | **p-value** |  |  | — | 0.033 | 0.056 |
| **SSF-LA** | **p-value** |  |  |  | — | 0.996 |
| **Untreated** | **p-value** |  |  |  |  | — |

| Tukey Post-Hoc Test – Guluronic acid | | | | | | |
| --- | --- | --- | --- | --- | --- | --- |
|  |  | **SMF-SP** | **SSF-SP** | **SMF-LA** | **SSF-LA** | **Untreated** |
| **SMF-SP** | **p-value** | — | 0.999 | 0.051 | 0.061 | 0.002 |
| **SSF-SP** | **p-value** |  | — | 0.036 | 0.086 | 0.002 |
| **SMF-LA** | **p-value** |  |  | — | <.001 | <.001 |
| **SSF-LA** | **p-value** |  |  |  | — | 0.166 |
| **Untreated** | **p-value** |  |  |  |  | — |

| Tukey Post-Hoc Test – Mannuronic acid | | | | | | |
| --- | --- | --- | --- | --- | --- | --- |
|  |  | **SMF-SP** | **SSF-SP** | **SMF-LA** | **SSF-LA** | **Untreated** |
| **SMF-SP** | **p-value** | — | 0.867 | 0.110 | 1.000 | 0.230 |
| **SSF-SP** | **p-value** |  | — | 0.415 | 0.915 | 0.055 |
| **SMF-LA** | **p-value** |  |  | — | 0.134 | 0.004 |
| **SSF-LA** | **p-value** |  |  |  | — | 0.193 |
| **Untreated** | **p-value** |  |  |  |  | — |

| Tukey Post-Hoc Test – Total Carb | | | | | | |
| --- | --- | --- | --- | --- | --- | --- |
|  |  | **SMF-SP** | **SSF-SP** | **SMF-LA** | **SSF-LA** | **Untreated** |
| **SMF-SP** | **p-value** | — | 0.872 | 0.149 | 0.614 | 0.019 |
| **SSF-SP** | **p-value** |  | — | 0.514 | 0.194 | 0.005 |
| **SMF-LA** | **p-value** |  |  | — | 0.016 | <.001 |
| **SSF-LA** | **p-value** |  |  |  | — | 0.173 |
| **Untreated** | **p-value** |  |  |  |  | — |

**Table S2. Statistics for protein analysis (*n* = 3).**

| One-Way ANOVA (Fisher's) | | | | |
| --- | --- | --- | --- | --- |
|  | **F** | **df1** | **df2** | **p** |
| **Protein** | 30.3 | 4 | 10 | <.001 |

| Tukey Post-Hoc Test – Protein | | | | | | |
| --- | --- | --- | --- | --- | --- | --- |
|  |  | **SMF-SP** | **SSF-SP** | **SMF-LA** | **SSF-LA** | **Untreated** |
| **SMF-SP** | **p-value** | — | 1.000 | 0.214 | 0.944 | <.001 |
| **SSF-SP** | **p-value** |  | — | 0.171 | 0.975 | <.001 |
| **SMF-LA** | **p-value** |  |  | — | 0.072 | <.001 |
| **SSF-LA** | **p-value** |  |  |  | — | <.001 |
| **Untreated** | **p-value** |  |  |  |  | — |

**Table S3.** **Statistics for total solids (TS%) and total mineral (Ash%) analysis (*n* = 3).**

| One-Way ANOVA (Fisher's) | | | | |
| --- | --- | --- | --- | --- |
|  | **F** | **df1** | **df2** | **p** |
| **TS%** | 0.672 | 3 | 8 | 0.593 |
| **Ash%** | 182.642 | 3 | 8 | <.001 |

| Tukey Post-Hoc Test – TS% | | | | | |
| --- | --- | --- | --- | --- | --- |
|  |  | **SMF-LA** | **SSF-LA** | **Untreated** | **Water soaked** |
| **SMF-LA** | **p-value** | — | 0.715 | 0.976 | 0.620 |
| **SSF-LA** | **p-value** |  | — | 0.907 | 0.998 |
| **Untreated** | **p-value** |  |  | — | 0.835 |
| **Water soaked** | **p-value** |  |  |  | — |

| Tukey Post-Hoc Test – Ash% | | | | | |
| --- | --- | --- | --- | --- | --- |
|  |  | **SMF-LA** | **SSF-LA** | **Untreated** | **Water soaked** |
| **SMF-LA** | **p-value** | — | 0.033 | <.001 | 1.000 |
| **SSF-LA** | **p-value** |  | — | <.001 | 0.038 |
| **Untreated** | **p-value** |  |  | — | <.001 |
| **Water soaked** | **p-value** |  |  |  | — |

**Table S4. Statistics for mineral analysis (*n* = 3).**

| One-Way ANOVA (Fisher's) | | | | |
| --- | --- | --- | --- | --- |
|  | **F** | **df1** | **df2** | **p** |
| **Ag** | 1.34 | 3 | 8 | 0.328 |
| **Ca** | 10.04 | 3 | 8 | 0.004 |
| **Co** | 36.70 | 3 | 8 | <.001 |
| **Cr** | 2.54 | 3 | 8 | 0.130 |
| **Cu** | 41.38 | 3 | 8 | <.001 |
| **Fe** | 36.62 | 3 | 8 | <.001 |
| **K** | 25.12 | 3 | 8 | <.001 |
| **Mg** | 65.19 | 3 | 8 | <.001 |
| **Mn** | 16.65 | 3 | 8 | <.001 |
| **Mo** | 8.16 | 3 | 8 | 0.008 |
| **Na** | 17.40 | 3 | 8 | <.001 |
| **Ni** | 7.22 | 3 | 8 | 0.012 |
| **P** | 3.19 | 3 | 8 | 0.084 |
| **Se** | 8.02 | 3 | 8 | 0.009 |
| **Zn** | 34.76 | 3 | 8 | <.001 |

| Tukey Post-Hoc Test – Ag | | | | | |
| --- | --- | --- | --- | --- | --- |
|  |  | **SMF-LA** | **SSF-LA** | **Untreated** | **Water soaked** |
| **SMF-LA** | **p-value** | — | 1.000 | 0.999 | 0.431 |
| **SSF-LA** | **p-value** |  | — | 0.997 | 0.458 |
| **Untreated** | **p-value** |  |  | — | 0.363 |
| **Water soaked** | **p-value** |  |  |  | — |

| Tukey Post-Hoc Test – Ca | | | | | |
| --- | --- | --- | --- | --- | --- |
|  |  | **SMF-LA** | **SSF-LA** | **Untreated** | **Water soaked** |
| **SMF-LA** | **p-value** | — | 0.425 | 0.006 | 1.000 |
| **SSF-LA** | **p-value** |  | — | 0.052 | 0.450 |
| **Untreated** | **p-value** |  |  | — | 0.006 |
| **Water soaked** | **p-value** |  |  |  | — |

| Tukey Post-Hoc Test – Co | | | | | |
| --- | --- | --- | --- | --- | --- |
|  |  | **SMF-LA** | **SSF-LA** | **Untreated** | **Water soaked** |
| **SMF-LA** | **p-value** | — | 0.467 | 0.009 | 0.002 |
| **SSF-LA** | **p-value** |  | — | 0.070 | <.001 |
| **Untreated** | **p-value** |  |  | — | <.001 |
| **Water soaked** | **p-value** |  |  |  | — |

| Tukey Post-Hoc Test – Cr | | | | | |
| --- | --- | --- | --- | --- | --- |
|  |  | **SMF-LA** | **SSF-LA** | **Untreated** | **Water soaked** |
| **SMF-LA** | **p-value** | — | 0.896 | 0.119 | 0.944 |
| **SSF-LA** | **p-value** |  | — | 0.303 | 0.999 |
| **Untreated** | **p-value** |  |  | — | 0.252 |
| **Water soaked** | **p-value** |  |  |  | — |

| Tukey Post-Hoc Test – Cu | | | | | |
| --- | --- | --- | --- | --- | --- |
|  |  | **SMF-LA** | **SSF-LA** | **Untreated** | **Water soaked** |
| **SMF-LA** | **p-value** | — | 0.114 | <.001 | 0.930 |
| **SSF-LA** | **p-value** |  | — | <.001 | 0.049 |
| **Untreated** | **p-value** |  |  | — | <.001 |
| **Water soaked** | **p-value** |  |  |  | — |

| Tukey Post-Hoc Test – Fe | | | | | |
| --- | --- | --- | --- | --- | --- |
|  |  | **SMF-LA** | **SSF-LA** | **Untreated** | **Water soaked** |
| **SMF-LA** | **p-value** | — | 0.743 | 0.002 | 0.009 |
| **SSF-LA** | **p-value** |  | — | 0.005 | 0.003 |
| **Untreated** | **p-value** |  |  | — | <.001 |
| **Water soaked** | **p-value** |  |  |  | — |

| Tukey Post-Hoc Test – K | | | | | |
| --- | --- | --- | --- | --- | --- |
|  |  | **SMF-LA** | **SSF-LA** | **Untreated** | **Water soaked** |
| **SMF-LA** | **p-value** | — | 0.031 | <.001 | 0.881 |
| **SSF-LA** | **p-value** |  | — | 0.011 | 0.088 |
| **Untreated** | **p-value** |  |  | — | <.001 |
| **Water soaked** | **p-value** |  |  |  | — |

| Tukey Post-Hoc Test – Mg | | | | | |
| --- | --- | --- | --- | --- | --- |
|  |  | **SMF-LA** | **SSF-LA** | **Untreated** | **Water soaked** |
| **SMF-LA** | **p-value** | — | 0.999 | <.001 | 0.818 |
| **SSF-LA** | **p-value** |  | — | <.001 | 0.877 |
| **Untreated** | **p-value** |  |  | — | <.001 |
| **Water soaked** | **p-value** |  |  |  | — |

| Tukey Post-Hoc Test – Mn | | | | | |
| --- | --- | --- | --- | --- | --- |
|  |  | **SMF-LA** | **SSF-LA** | **Untreated** | **Water soaked** |
| **SMF-LA** | **p-value** | — | 0.036 | <.001 | 0.004 |
| **SSF-LA** | **p-value** |  | — | 0.042 | 0.415 |
| **Untreated** | **p-value** |  |  | — | 0.385 |
| **Water soaked** | **p-value** |  |  |  | — |

| Tukey Post-Hoc Test – Mo | | | | | |
| --- | --- | --- | --- | --- | --- |
|  |  | **SMF-LA** | **SSF-LA** | **Untreated** | **Water soaked** |
| **SMF-LA** | **p-value** | — | 0.443 | 0.009 | 0.027 |
| **SSF-LA** | **p-value** |  | — | 0.079 | 0.237 |
| **Untreated** | **p-value** |  |  | — | 0.853 |
| **Water soaked** | **p-value** |  |  |  | — |

| Tukey Post-Hoc Test – Na | | | | | |
| --- | --- | --- | --- | --- | --- |
|  |  | **SMF-LA** | **SSF-LA** | **Untreated** | **Water soaked** |
| **SMF-LA** | **p-value** | — | 0.251 | 0.002 | 0.789 |
| **SSF-LA** | **p-value** |  | — | 0.026 | 0.069 |
| **Untreated** | **p-value** |  |  | — | <.001 |
| **Water soaked** | **p-value** |  |  |  | — |

| Tukey Post-Hoc Test – Ni | | | | | |
| --- | --- | --- | --- | --- | --- |
|  |  | **SMF-LA** | **SSF-LA** | **Untreated** | **Water soaked** |
| **SMF-LA** | **p-value** | — | 0.117 | 0.008 | 0.392 |
| **SSF-LA** | **p-value** |  | — | 0.278 | 0.796 |
| **Untreated** | **p-value** |  |  | — | 0.079 |
| **Water soaked** | **p-value** |  |  |  | — |

| Tukey Post-Hoc Test – P | | | | | |
| --- | --- | --- | --- | --- | --- |
|  |  | **SMF-LA** | **SSF-LA** | **Untreated** | **Water soaked** |
| **SMF-LA** | **p-value** | — | 0.846 | 0.350 | 0.642 |
| **SSF-LA** | **p-value** |  | — | 0.775 | 0.258 |
| **Untreated** | **p-value** |  |  | — | 0.069 |
| **Water soaked** | **p-value** |  |  |  | — |

| Tukey Post-Hoc Test – Se | | | | | |
| --- | --- | --- | --- | --- | --- |
|  |  | **SMF-LA** | **SSF-LA** | **Untreated** | **Water soaked** |
| **SMF-LA** | **p-value** | — | 0.158 | 0.010 | 0.975 |
| **SSF-LA** | **p-value** |  | — | 0.274 | 0.274 |
| **Untreated** | **p-value** |  |  | — | 0.018 |
| **Water soaked** | **p-value** |  |  |  | — |

| Tukey Post-Hoc Test – Zn | | | | | |
| --- | --- | --- | --- | --- | --- |
|  |  | **SMF-LA** | **SSF-LA** | **Untreated** | **Water soaked** |
| **SMF-LA** | **p-value** | — | 0.518 | 0.075 | <.001 |
| **SSF-LA** | **p-value** |  | — | 0.485 | <.001 |
| **Untreated** | **p-value** |  |  | — | <.001 |
| **Water soaked** | **p-value** |  |  |  | — |

**Table S5. Statistics for potentially toxic element (PTE) analysis (*n* = 3).**

| One-Way ANOVA (Fisher's) | | | | |
| --- | --- | --- | --- | --- |
|  | **F** | **df1** | **df2** | **p** |
| **I** | 97.15 | 3 | 8 | <.001 |
| **As** | 6.73 | 3 | 8 | 0.014 |
| **Cd** | 25.05 | 3 | 8 | <.001 |
| **Pb** | 9.52 | 3 | 8 | 0.005 |

| Tukey Post-Hoc Test – I | | | | | |
| --- | --- | --- | --- | --- | --- |
|  |  | **SMF-LA** | **SSF-LA** | **Untreated** | **Water soaked** |
| **SMF-LA** | **p-value** | — | 0.265 | 0.002 | <.001 |
| **SSF-LA** | **p-value** |  | — | 0.023 | <.001 |
| **Untreated** | **p-value** |  |  | — | <.001 |
| **Water soaked** | **p-value** |  |  |  | — |

| Tukey Post-Hoc Test – As | | | | | |
| --- | --- | --- | --- | --- | --- |
|  |  | **SMF-LA** | **SSF-LA** | **Untreated** | **Water soaked** |
| **SMF-LA** | **p-value** | — | 0.597 | 0.012 | 0.733 |
| **SSF-LA** | **p-value** |  | — | 0.068 | 0.995 |
| **Untreated** | **p-value** |  |  | — | 0.048 |
| **Water soaked** | **p-value** |  |  |  | — |

| Tukey Post-Hoc Test – Cd | | | | | |
| --- | --- | --- | --- | --- | --- |
|  |  | **SMF-LA** | **SSF-LA** | **Untreated** | **Water soaked** |
| **SMF-LA** | **p-value** | — | 0.998 | 0.168 | 0.001 |
| **SSF-LA** | **p-value** |  | — | 0.134 | 0.002 |
| **Untreated** | **p-value** |  |  | — | <.001 |
| **Water soaked** | **p-value** |  |  |  | — |

| Tukey Post-Hoc Test – Pb | | | | | |
| --- | --- | --- | --- | --- | --- |
|  |  | **SMF-LA** | **SSF-LA** | **Untreated** | **Water soaked** |
| **SMF-LA** | **p-value** | — | 0.683 | 0.072 | 0.237 |
| **SSF-LA** | **p-value** |  | — | 0.015 | 0.778 |
| **Untreated** | **p-value** |  |  | — | 0.004 |
| **Water soaked** | **p-value** |  |  |  | — |

**Table S6.** Composition of untreated *Alaria esculenta.*

| **Components** | **Amount** |
| --- | --- |
| Total solids (g/100g DW) | 91.58 ± 0.09 |
| Total minerals (ash) (g/100g DW) | 37.57 ± 0.60 |
| Protein (% DW) | 11.6 ± 0.30 |
| Total carbohydrate (g/100g DW) | 16.57 ± 1.40 |
| Mannitol | 5.40 ± 0.10 |
| Fucose | 0.91 ± 0.03 |
| Glucose | 1.88 ± 0.16 |
| Xylose | 0.22 ± 0.04 |
| Mannose | 0.03 ± 0.03 |
| Glucuronic acid | 1.67 ± 0.610 |
| Guluronic acid | 1.03 ± 0.07 |
| Mannuronic acid | 5.44 ± 0.97 |
| Mineral content (mg/kg DW) |  |
| K | 79 300 ± 2 443 |
| Na | 48 167 ± 1 966 |
| Ca | 11 533 ± 723 |
| Mg | 11 567 ± 493 |
| P | 3 817 ± 146 |
| Fe | 127 ± 6 |
| Zn | 51 ± 2 |
| Mn | 6.90 ± 0.09 |
| Cu | 3.29 ± 0.48 |
| Ni | 1.75 ± 0.06 |
| Cr | 1.36 ± 0.18 |
| Co | 0.123 ± 0.005 |
| Se | 0.306 ± 0.015 |
| Mo | 0.205 ± 0.004 |
| Ag | 0.020 ± 0.002 |
| Potentially toxic elements (mg/kg DW) |  |
| I | 808 ± 35 |
| As | 50.7 ± 0.5 |
| Cd | 1.37 ± 0.06 |
| Pb | 0.156 ± 0.003 |
| Hg | < 0.020 |
| Antioxidant compounds (mg/g DW) |  |
| Phloroglucinol | 0.68±0.10 |
| Epicatechin | 0.0061±0.001 |
| Salicylic acid | 0.033±0.01 |
| Syringic acid | <0.0008 |
| Ferrulic acid | 0.0014±0.0001 |
| 3,4-DHBA | 0.0050±0.0005 |
| 4-HBA | 0.0030±0.0005 |
| Vanilic acid | 0.0057±0.0008 |
| Caffeic acid | <0.0005 |
| Chlorogenic acid | <0.0006 |
| Rutin | <0.0007 |
| Quercetin | <0.0007 |
| Kaempferol | 0.0021±0.0004 |
| β-Carotene | 0.24±0.006 |
| Fucoxanthin | 0.87±0.05 |
| α-Tocopherol | 0.018±0.001 |

**Table S7.** Antioxidant composition of untreated Alaria esculenta and of both the solid residue and supernatant from seaweed fermented under submerged conditions (SmF-LA and SmF-SP). Variables marked with an asterisk (*) have *p*-values greater than 0.05, indicating no significant difference.

| Antioxidant compound | mg/g DW | | | | | |
| --- | --- | --- | --- | --- | --- | --- |
|  | **Untreated** | **SmF-SP**  **(Residue)** | **SmF-LA (Residue)** | **SmF-SP (Supernatant)** | **SmF-LA (Supernatant)** | ***p*-value**  **(ANOVA)** |
| Phloroglucinol | 0.68±0.10 | 1.49±0.02 | 2.06±0.5 | 2.61±0.4 | 4.06±0.3 | <0.001 |
| Epicatechin | 0.0061±0.001 | 0.0017±0.00001 | 0.0035±0.0006 | 0.0049±0.001 | 0.0073±0.001 | 0.003 |
| Salicylic acid | 0.033±0.01 | 0.022±0.00003 | 0.011±0.003 | 0.010±0.001 | 0.011±0.002 | <0.001 |
| Syringic acid | <0.0008 | <0.0008 | 0.0021±0.0006 | 0.0037±0.0004 | 0.0064±0.0002 | <0.001 |
| Ferulic acid | 0.0014±0.0001 | 0.00091±0.00001 | 0.00085±0.0001 | 0.0038±0.0002 | 0.0018±0.0002 | <0.001 |
| 3,4-DHBA | 0.0050±0.0005 | 0.0044±0.0007 | <0.0013 | <0.0013 | 0.0032±0.0005 | 0.003 |
| 4-HBA | 0.0030±0.0005 | 0.0046±0.0007 | 0.0026±0.0003 | 0.0026±0.0002 | 0.0028±0.0002 | 0.001 |
| Vanillic acid | 0.0057±0.0008 | <0.0014 | <0.0014 | 0.0050±0.0001 | <0.0014 | 0.14 * |
| Caffeic acid | <0.0005 | 0.0005±0.0001 | 0.0006±0.0002 | 0.0012±0.00007 | 0.0015±0.00008 | <0.001 |
| Chlorogenic acid | <0.0006 | 0.0010±0.0002 | 0.0010± | 0.0021±0.00002 | 0.0020±0.0001 | <0.001 |
| Rutin | <0.0007 | <0.0007 | 0.0007± | 0.0074±0.00071 | <0.0007 | - |
| Quercetin | <0.0007 | 0.0016±0.0003 | 0.0013±0.0001 | <0.0007 | <0.0007 | 0.13* |
| Kaempferol | 0.0021±0.0004 | 0.0027±0.0003 | 0.0018±0.00009 | <0.0008 | <0.0008 | 0.08* |
| β-Carotene | 0.24±0.006 | 0.062±0.012 | 0.028±0.01 | <0.0023 | <0.0023 | <0.001 |
| Fucoxanthin | 0.87±0.05 | 0.34±0.07 | 0.045±0.007 | <0.0024 | <0.0024 | <0.001 |
| α-Tocopherol | 0.018±0.001 | 0.016±0.001 | 0.013±0.006 | <0.0007 | <0.0007 | 0.45* |

**Table S8.** Tukey’s HSD test results for antioxidant compounds that showed significant differences between treatments in the ANOVA analysis.

| Tukey Post-Hoc Test – Phloroglucinol | | | | | | |
| --- | --- | --- | --- | --- | --- | --- |
|  |  | **SMF-SP (Residue)** | **SMF-LA (Residue)** | **SMF-SP (Supernatant)** | **SMF-LA (Supernatant)** | **Untreated** |
| **SMF-SP (Residue)** | **p-value** | — | 0.383 | 0.28 | <0.001 | 0.126 |
| **SMF-LA (Residue)** | **p-value** |  | — | 0.304 | <0.001 | 0.004 |
| **SMF-SP (Supernatant)** | **p-value** |  |  | — | 0.003 | <0.001 |
| **SMF-LA (Supernatant)** | **p-value** |  |  |  | — | <.001 |
| **Untreated** | **p-value** |  |  |  |  | — |

| Tukey Post-Hoc Test – Epicatechin | | | | | | |
| --- | --- | --- | --- | --- | --- | --- |
|  |  | **SMF-SP (Residue)** | **SMF-LA (Residue)** | **SMF-SP (Supernatant)** | **SMF-LA (Supernatant)** | **Untreated** |
| **SMF-SP (Residue)** | **p-value** | — | 0.452 | 0.069 | 0.003 | 0.015 |
| **SMF-LA (Residue)** | **p-value** |  | — | 0.557 | 0.018 | 0.12 |
| **SMF-SP (Supernatant)** | **p-value** |  |  | — | 0.173 | 0.749 |
| **SMF-LA (Supernatant)** | **p-value** |  |  |  | — | 0.7 |
| **Untreated** | **p-value** |  |  |  |  | — |

| Tukey Post-Hoc Test – Salicylic acid | | | | | | |
| --- | --- | --- | --- | --- | --- | --- |
|  |  | **SMF-SP (Residue)** | **SMF-LA (Residue)** | **SMF-SP (Supernatant)** | **SMF-LA (Supernatant)** | **Untreated** |
| **SMF-SP (Residue)** | **p-value** | — | 0.326 | 0.272 | 0.322 | 0.244 |
| **SMF-LA (Residue)** | **p-value** |  | — | 1.00 | 1.00 | 0.007 |
| **SMF-SP (Supernatant)** | **p-value** |  |  | — | 1.00 | 0.005 |
| **SMF-LA (Supernatant)** | **p-value** |  |  |  | — | 0.007 |
| **Untreated** | **p-value** |  |  |  |  | — |

| Tukey Post-Hoc Test – 4-HBA | | | | | | |
| --- | --- | --- | --- | --- | --- | --- |
|  |  | **SMF-SP (Residue)** | **SMF-LA (Residue)** | **SMF-SP (Supernatant)** | **SMF-LA (Supernatant)** | **Untreated** |
| **SMF-SP (Residue)** | **p-value** | — | 0.002 | 0.002 | 0.005 | 0.011 |
| **SMF-LA (Residue)** | **p-value** |  | — | 1.00 | 0.943 | 0.695 |
| **SMF-SP (Supernatant)** | **p-value** |  |  | — | 0.976 | 0.781 |
| **SMF-LA (Supernatant)** | **p-value** |  |  |  | — | 0.977 |
| **Untreated** | **p-value** |  |  |  |  | — |

| Tukey Post-Hoc Test – Ferulic acid | | | | | | |
| --- | --- | --- | --- | --- | --- | --- |
|  |  | **SMF-SP (Residue)** | **SMF-LA (Residue)** | **SMF-SP (Supernatant)** | **SMF-LA (Supernatant)** | **Untreated** |
| **SMF-SP (Residue)** | **p-value** | — | 0.993 | < 0.001 | < 0.001 | 0.022 |
| **SMF-LA (Residue)** | **p-value** |  | — | < 0.001 | < 0.001 | 0.006 |
| **SMF-SP (Supernatant)** | **p-value** |  |  | — | < 0.001 | < 0.001 |
| **SMF-LA (Supernatant)** | **p-value** |  |  |  | — | 0.037 |
| **Untreated** | **p-value** |  |  |  |  | — |

| Tukey Post-Hoc Test – Syringic acid | | | | | | |
| --- | --- | --- | --- | --- | --- | --- |
|  |  | **SMF-SP (Residue)** | **SMF-LA (Residue)** | **SMF-SP (Supernatant)** | **SMF-LA (Supernatant)** | **Untreated** |
| **SMF-SP (Residue)** | **p-value** | — | N/A | N/A | N/A | N/A |
| **SMF-LA (Residue)** | **p-value** |  | — | 0.01 | < 0.001 | N/A |
| **SMF-SP (Supernatant)** | **p-value** |  |  | — | < 0.001 | N/A |
| **SMF-LA (Supernatant)** | **p-value** |  |  |  | — | N/A |
| **Untreated** | **p-value** |  |  |  |  | — |

| Tukey Post-Hoc Test – 3, 4-DHBA | | | | | | |
| --- | --- | --- | --- | --- | --- | --- |
|  |  | **SMF-SP (Residue)** | **SMF-LA (Residue)** | **SMF-SP (Supernatant)** | **SMF-LA (Supernatant)** | **Untreated** |
| **SMF-SP (Residue)** | **p-value** | — | N/A | N/A | 0.076 | 0.05 |
| **SMF-LA (Residue)** | **p-value** |  | — | N/A | N/A | N/A |
| **SMF-SP (Supernatant)** | **p-value** |  |  | — | N/A | N/A |
| **SMF-LA (Supernatant)** | **p-value** |  |  |  | — | 0.003 |
| **Untreated** | **p-value** |  |  |  |  | — |

| Tukey Post-Hoc Test – Caffeic acid | | | | | | |
| --- | --- | --- | --- | --- | --- | --- |
|  |  | **SMF-SP (Residue)** | **SMF-LA (Residue)** | **SMF-SP (Supernatant)** | **SMF-LA (Supernatant)** | **Untreated** |
| **SMF-SP (Residue)** | **p-value** | — | 0.861 | 0.001 | < 0.001 | N/A |
| **SMF-LA (Residue)** | **p-value** |  | — | 0.003 | < 0.001 | N/A |
| **SMF-SP (Supernatant)** | **p-value** |  |  | — | 0.27 | N/A |
| **SMF-LA (Supernatant)** | **p-value** |  |  |  | — | N/A |
| **Untreated** | **p-value** |  |  |  |  | — |

| Tukey Post-Hoc Test – Chlorogenic acid | | | | | | |
| --- | --- | --- | --- | --- | --- | --- |
|  |  | **SMF-SP (Residue)** | **SMF-LA (Residue)** | **SMF-SP (Supernatant)** | **SMF-LA (Supernatant)** | **Untreated** |
| **SMF-SP (Residue)** | **p-value** | — | N/A | < 0.001 | < 0.001 | N/A |
| **SMF-LA (Residue)** | **p-value** |  | — | N/A | N/A | N/A |
| **SMF-SP (Supernatant)** | **p-value** |  |  | — | 0.778 | N/A |
| **SMF-LA (Supernatant)** | **p-value** |  |  |  | — | N/A |
| **Untreated** | **p-value** |  |  |  |  | — |

| Tukey Post-Hoc Test – β-Carotene | | | | | | |
| --- | --- | --- | --- | --- | --- | --- |
|  |  | **SMF-SP (Residue)** | **SMF-LA (Residue)** | **Untreated** | **SMF-LA (Supernatant)** | **SMF-SP (Supernatant)** |
| **SMF-SP (Residue)** | **p-value** | — | 0.022 | < 0.001 | N/A | N/A |
| **SMF-LA (Residue)** | **p-value** |  | — | < 0.001 | N/A | N/A |
| **Untreated** | **p-value** |  |  | — | N/A | N/A |
| **SMF-LA (Supernatant)** | **p-value** |  |  |  | — | N/A |
| **SMF-SP (Supernatant)** | **p-value** |  |  |  |  | — |

| Tukey Post-Hoc Test – Fucoxanthin | | | | | | |
| --- | --- | --- | --- | --- | --- | --- |
|  |  | **SMF-SP (Residue)** | **SMF-LA (Residue)** | **Untreated** | **SMF-LA (Supernatant)** | **SMF-SP (Supernatant)** |
| **SMF-SP (Residue)** | **p-value** | — | < 0.001 | < 0.001 | N/A | N/A |
| **SMF-LA (Residue)** | **p-value** |  | — | < 0.001 | N/A | N/A |
| **Untreated** | **p-value** |  |  | — | N/A | N/A |
| **SMF-LA (Supernatant)** | **p-value** |  |  |  | — | N/A |
| **SMF-SP (Supernatant)** | **p-value** |  |  |  |  | — |

**NB:** N/A indicates treatments in which concentrations were below the limit of quantification (LOQ) and therefore not applicable.
